# An Evidence-Grounded Research Assistant for Functional Genomics and Drug Target Assessment

**DOI:** 10.64898/2025.12.30.697073

**Authors:** Ksenia Sokolova, Dmitri Kosenkov, Keerthana Nallamotu, Sanketh Vedula, Daniil Sokolov, Guillermo Sapiro, Olga G Troyanskaya

**Affiliations:** Princeton Precision Health, Princeton University, Princeton, NJ, USA; Flatiron Institute, Simons Foundation, New York City, NY, USA; Department of Computer Science, Princeton University, Princeton, NJ, USA; Lewis-Sigler Institute for Integrative Genomics, Princeton University, Princeton, NJ, USA; Department of Electrical and Computer Engineering, Princeton University, Princeton, NJ, USA; Apple, Cupertino, CA, USA

## Abstract

The growing availability of biological data resources has transformed research, yet their effective use remains challenging: selecting appropriate sources requires domain knowledge, data are fragmented across databases, and synthesizing results into reliable conclusions is labor-intensive. Although large language models promise to address these barriers, their impact in biomedicine has been limited by unsupported statements, incorrect claims, and lack of provenance. We introduce Alvessa, an evidence-grounded agentic research assistant designed around verifiability. Alvessa integrates entity recognition, orchestration of pre-validated biological tools, and data-constrained answer generation with statement-level verification against retrieved records, explicitly flagging unsupported claims and guiding revision when reliability criteria are not met. We evaluate Alvessa on dbQA from LAB-Bench and GenomeArena, a benchmark of 720 questions spanning gene and variant annotation, pathways, molecular interactions, miRNA targets, drug-target evidence, protein structure, and gene-phenotype associations. Alvessa substantially improves accuracy relative to general-purpose language models and performs comparably to coding-centric agents while producing fully traceable outputs. Using adversarial perturbations, we show that detection of fabricated statements depends critically on access to retrieved evidence. We further demonstrate application to drug discovery, where evidence-grounded synthesis enables identification of candidate targets missed or misattributed by literature-centered reasoning alone. Alvessa and GenomeArena are released to the community to support reproducible, verifiable AI-assisted biological research.

## Introduction

The growing availability of diverse data resources, including variant annotations, pathway knowledge, interaction data, and chemoproteomics profiles, has transformed biological research. Yet using these resources remains challenging: selecting appropriate sources requires domain expertise, and the data are fragmented across databases with incompatible identifiers, schemas, and update cycles. Even seemingly straightforward questions, such as mapping a reported variant to the correct coordinate system, linking it to a gene and phenotype, and reconciling evidence across sources, often require significant time and technical expertise. Once data is collected, synthesizing it into coherent conclusions remains a separate, often equally demanding task.

Large language models (LLMs) offer an appealing solution, but they routinely produce answers that are overconfident, incorrect, or untraceable. The problem is amplified for genomics: identifier spaces are vast, fine-grained distinctions carry meaning (rsIDs, Ensembl accessions, transcript isoforms), and plausible-sounding errors can be difficult to detect before propagating into downstream analyses. For example, dbSNP alone now catalogs over 1.2 billion reference SNP identifiers (“rsIDs”)^1^, making LLM memorization and reliable disambiguation unrealistic. These limitations highlight a fundamental challenge for scientific AI: correctness and provenance must be enforced at the level of individual claims, not inferred from fluent generation.

Recent agentic systems extend LLMs with planning and tool use^2–5^. For example, leading LLMs now use web search to access available information or perform literature search over published articles. Yet provenance is rarely enforced at the level required for scientific reliability; users may see links or high-level citations, but individual output statements are not systematically traced to retrieved records, as expected from good human and machine research assistants. As a result, these systems can produce outputs that appear reasonable, yet contain fabricated identifiers, incorrect numeric values, or claims only partially supported by evidence.

Coding agents, which dynamically generate and execute code, offer a complementary approach. Recent systems such as Biomni^6^ use agentic workflows to analyze data on the fly with access to a variety of bioinformatic tools and complementary data sources. However, these systems are primarily designed to analyze user-provided data and execute procedural workflows, rather than to surface specific records with systematic, statement-level provenance. In addition, on-the-fly code generation can lead to variability in analyses and outputs across runs, complicating reproducibility and interpretation.

To address these challenges, in particular for genomics, here we introduce Alvessa, a multi-agent framework that acts as a reliable research assistant and treats verification as a core component of genomic reasoning. Alvessa operates by interpreting the user’s intent and dynamically orchestrating a catalog of validated tools to recognize entities and retrieve relevant data, generating answers that are strictly constrained to these records. Crucially, the answer is reviewed by a dedicated verification agent to scrutinize every output against the retrieved evidence, providing per-statement feedback and triggering re-writing of the answer if needed. To evaluate the basic knowledge required for reliable genomic assistance, we introduce GenomeArena, a curated benchmark of 720 questions systematically spanning variants, genes, pathways, interactions, regulatory targets, druggability, structure, and gene-phenotype associations; we also evaluate Alvessa on dbQA from LAB-Bench. We further stress-test verification with adversarial perturbations that inject plausible contradictions and identifier/numeric corruptions into otherwise grounded answers, directly testing whether the framework is capable of detecting these issues.

Finally, we illustrate how Alvessa can be used in the drug discovery pipeline. By synthesizing fragmented evidence across chemoproteomics, structural biology, and pharmacology, the system supports evidence-grounded assessment of high-confidence covalent ligandability opportunities within the ActRII pathway that standard literature-based reasoning obscures.

The proposed framework demonstrates that by enforcing strict alignment between generation and evidence, and providing reliable tools, language model agents can go beyond literature summarization to become reliable research assistants in fields where accuracy is critical. While demonstrated for biological questions, such systems can be extended to other areas of scientific discovery to act as collaborators of human scientists.

## Results

Alvessa is an agentic research assistant designed for text-based biomedical queries, built around the principle of trustworthiness through evidence provenance and systematic verification. The system comprises four main components: a purpose-built entity recognition module, intent understanding with tool orchestration, evidence-driven answer generation, and multi-step verification (Figure 1A). In the results below, we evaluate these capabilities by assessing (i) database routing and evidence retrieval on multiple-choice benchmarks and (ii) statement-level verification under adversarial perturbations.

**Figure 1.**
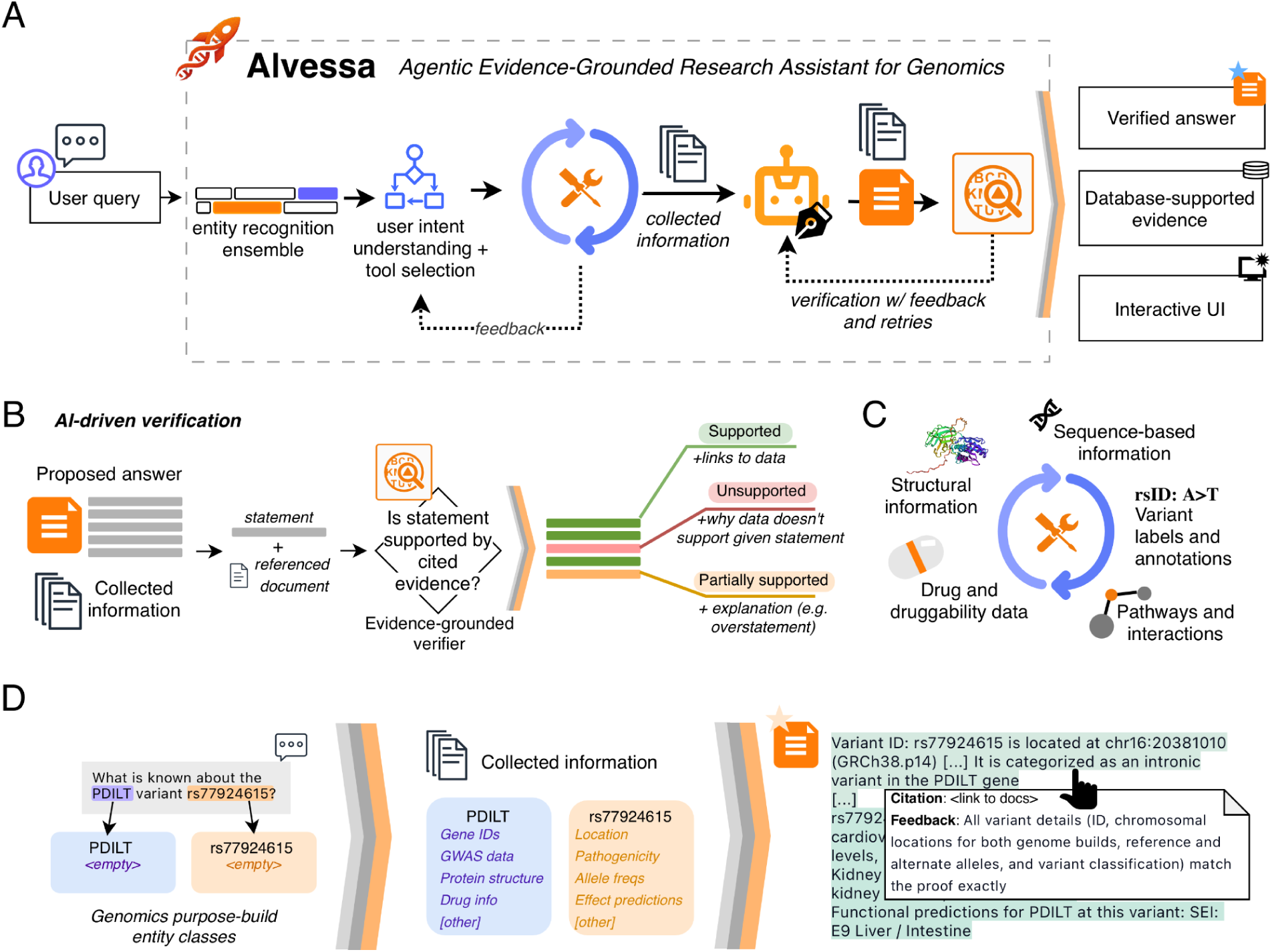
Alvessa architecture and workflow. **(A)** Overview of the Alvessa pipeline. User queries pass through entity recognition, intent understanding and tool selection, evidence collection, answer generation, and verification. Collected information is used to generate answers, with each statement having direct citations (references). Answers that fail verification return to the generation module with feedback. **(B)** The verification agent evaluates each statement against its cited evidence, classifying it as supported, unsupported, or partially supported, with explanations for non-supported statements. Global opinion about the answer is also generated. **(C)** Overview of the broad categories of information supported by the core set of Alvessa tools. **(D)** Schematic example of the workflow showing entity instantiation from a user query, population of structured information for each entity, and example view of the verified output with citation links and verification feedback.

When a user submits a query in natural language (much like a professor will ask their research assistant, or a PhD student might need to investigate as part of their dissertation), Alvessa identifies relevant entities, instantiates them as structured objects, and invokes appropriate tools to collect information spanning variant annotations, structural and sequence-based information, pathways and interactions, and drug and druggability data (Figure 1C-D, Methods). The system can expand its search by invoking additional tools if needed. Critically, all tools are pre-defined and validated rather than created on-the-fly, ensuring consistent and reliable behavior across runs.

The collected evidence is then used by the writing agent to generate a data-driven answer. For free-form responses, Alvessa then applies a verification agent that evaluates each statement against its cited evidence, classifying it as supported, partially supported, or unsupported (Figure 1B). Answers that fail verification return to the generation module with specific feedback for revision; when no relevant information is available, Alvessa reports this directly rather than generating unsupported claims.

Verified answers are presented in an interface where each statement displays its evidence linkage and reliability assessment, with underlying data available for download.

### Accurate performance across broad question categories

To evaluate whether Alvessa can reliably answer the base queries that underpin complex genomic reasoning, we developed GenomeArena, an open benchmark of 720 multiple-choice questions spanning eight categories: variant annotation, gene annotation, pathways, interactions, miRNA targets, drug-target relationships, protein structure, and gene-phenotype associations (Figure 2A). Questions were designed to both assess appropriate resource selection and factual accuracy; questions omit explicit database mentions unless needed for a deterministic setup, testing both factual accuracy and appropriate resource selection. To isolate database routing and evidence retrieval performance, we evaluate Alvessa on GenomeArena in a multiple-choice setting with verification disabled. The primary goal of this benchmark is to evaluate whether an agent can reliably route queries to the correct genomic resource and extract grounded answers, a prerequisite for downstream scientific reasoning.

**Figure 2.**
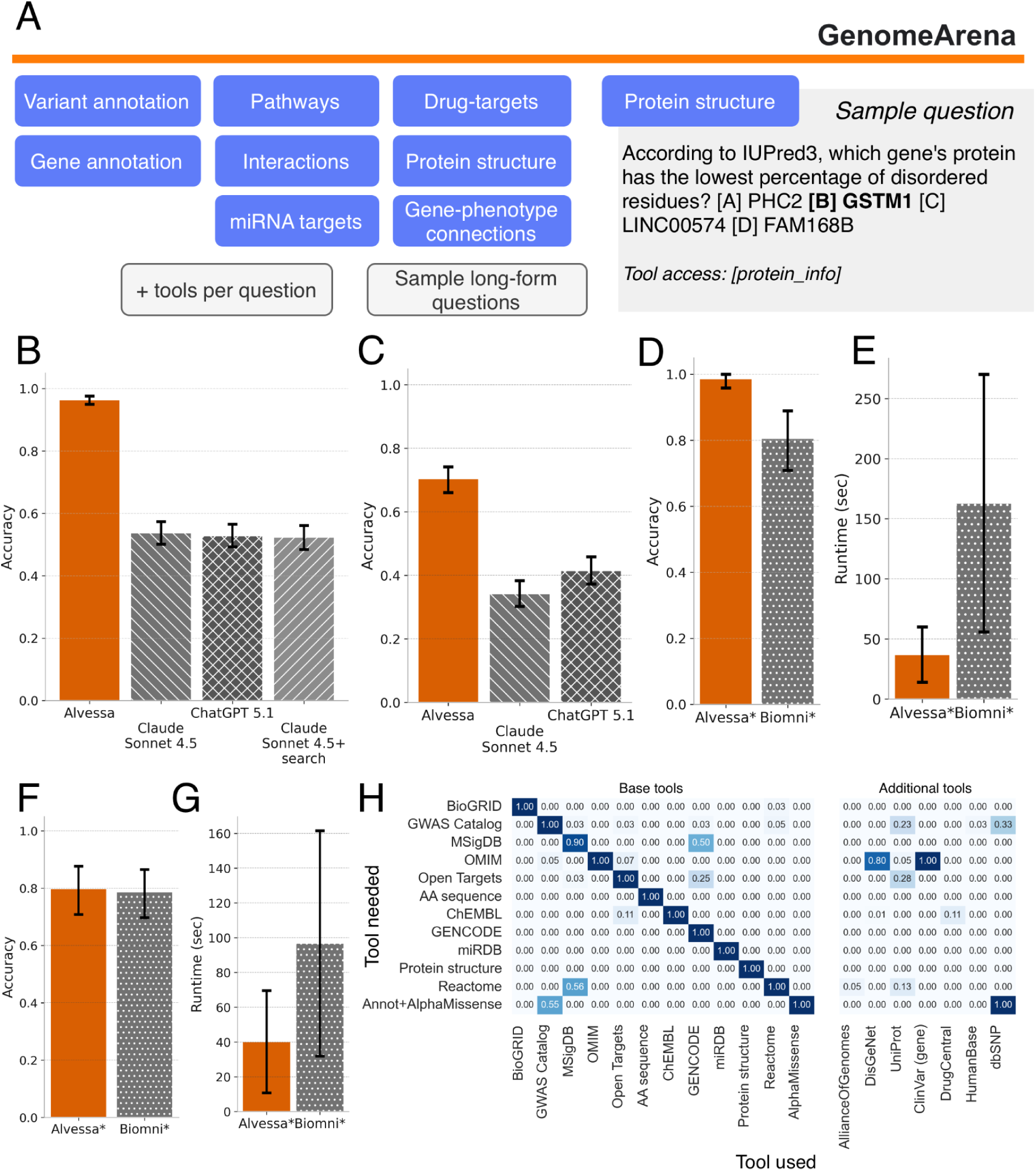
GenomeArena benchmark and evaluation. **(A)** GenomeArena comprises 720 multiple-choice questions spanning eight categories, each annotated with the expected primary tool. Sample long-form questions used for downstream verification analysis are also provided. **(B-D, F)** Accuracy scores across different benchmarks and models; error bars represent 95% bootstrapped confidence intervals. **(B)** Accuracy on GenomeArena comparing Alvessa to Claude Sonnet 4.5, ChatGPT 5.1, and Claude Sonnet 4.5 with integrated web search. **(C)** Accuracy on dbQA (DisGeNet questions excluded; see Methods). **(D)** Accuracy on a randomly sampled GenomeArena subset (n=72) comparing Alvessa and Biomni. **(E)** Mean time per question for Alvessa and Biomni on the GenomeArena subset; error bars indicate standard deviation. **(F)** Accuracy on a dbQA subset (n=89) comparing Alvessa and Biomni. **(G)** Mean time per question on the dbQA subset. **(H)** Tool selection matrix. Rows indicate the expected tool based on question metadata; columns indicate tools called. Diagonal values reflect correct primary tool selection; off-diagonal values indicate supplementary tools invoked (e.g., OMIM questions appropriately triggering ClinVar and DisGeNet).

Alvessa achieved 0.964 (95% CI: 0.949-0.976) accuracy on GenomeArena, substantially outperforming both Claude Sonnet 4.5 (0.537; 95% CI: 0.501-0.576) and ChatGPT 5.1 (0.528; 95% CI: 0.492-0.565) (Figure 2B). To test whether general-purpose information access could close this gap, we evaluated Claude Sonnet 4.5 with its integrated web search, allowing the model to autonomously formulate and execute queries as a researcher typically would. Web search did not improve overall performance, achieving accuracy of 0.524 (95% CI: 0.489-0.561). In particular, while disease-to-gene associations showed modest gains, protein-and miRNA-specific queries saw negligible benefit, and in some cases web search introduced noise that degraded accuracy (Supplementary Figure 2). These results indicate that access to information alone is insufficient, and structured retrieval from validated sources is important for reliable genomic question-answering.

We additionally evaluated Alvessa on dbQA from the LabBench suite^7^, an independent benchmark designed to test model performance across biological databases. After removing questions with annotation errors (see Methods for details), Alvessa achieved 0.704 (95% CI: 0.660-0.742) accuracy compared to 0.342 (95% CI: 0.302-0.383) for Claude Sonnet 4.5 and 0.415 (95% CI: 0.373-0.458) for ChatGPT 5.1 (Figure 2C).

Finally, we compared Alvessa to Biomni, a recently developed coding agent that generates and executes code dynamically. Due to Biomni’s computational cost (mean of 163 seconds per question), this comparison was performed on a randomly sampled subset of GenomeArena (10% of questions, n=72) and dbQA (n=89). On GenomeArena, Alvessa achieved 0.986 (95% CI: 0.958-1.000) accuracy compared to 0.806 (95% CI: 0.708-0.889) for Biomni; on dbQA, performance was comparable (Alvessa = 0.798, 95% CI: 0.708-0.876; Biomni=0.787, 95% CI: 0.697-0.865) (Figure 2D,F). Notably, Alvessa completed queries up to 4 times faster (mean time per question: 40 sec for Alvessa vs. 97 sec for Biomni on dbQA; 37 sec vs. 163 sec on GenomeArena; Figure 2E,G), a feature critical for scalability, while providing directly traceable tool calls and data provenance rather than dynamically generated code; these are key properties for a reliable research assistant. Although accuracy differences may partly reflect differences in tool coverage, these results demonstrate that pre-validated tool orchestration can match or exceed code-generation approaches while offering transparency and efficiency.

### Alvessa identifies appropriate entities and tools from query context

Reliable performance depends on accurate entity recognition, as missed or incorrectly resolved entities cannot be queried and directly limit downstream evidence retrieval. Recall was high across all evaluated entity types (Supplementary Figure 1 and Supplementary Table 3). Drug entities, the most challenging category due to naming variability, are handled through fuzzy matching against a comprehensive library to prioritize recall; downstream agents subsequently use only relevant information. For protein sequences, Alvessa employs a two-stage resolution strategy: exact substring matching followed by approximate k-mer search over a precomputed index, enabling rapid identification of both full-length and partial sequences without external alignment services.

Beyond entity recognition, Alvessa reliably selects the appropriate primary tool for almost all of the question categories (Figure 2H, Methods). The system also invokes complementary tools when relevant; for example, for miRDB questions, for OMIM-based disease queries the OMIM tool is correctly triggered while additionally calling ClinVar and DisGeNet, reflecting awareness that multiple databases may contain supporting evidence for gene-phenotype relationships. Questions requiring variant annotation coupled with AlphaMissense predictions correctly query AlphaMissense and use dbSNP or GWAS catalog to assign variants to genes.

### Alvessa detects unsupported statements through evidence grounding

To evaluate the robustness of Alvessa’s verification system, we developed an evaluation framework in which an adversarial LLM-based agent introduces errors into some of the statements within the generated answer before verification (Figure 3A, Supplementary Table 4, Methods). To evaluate over different questions, we use sample long-form questions from GenomeArena. When generating a false statement, the adversarial agent receives the original statement and has access to the cited evidence in context, enabling it to craft plausible but incorrect statements. We tested four error types: direct contradictions to cited evidence, overstatements that exaggerate beyond what evidence supports, wrong numerical values, and wrong alphanumeric identifiers such as replacing rsIDs or gene IDs (Figure 3B).

**Figure 3.**
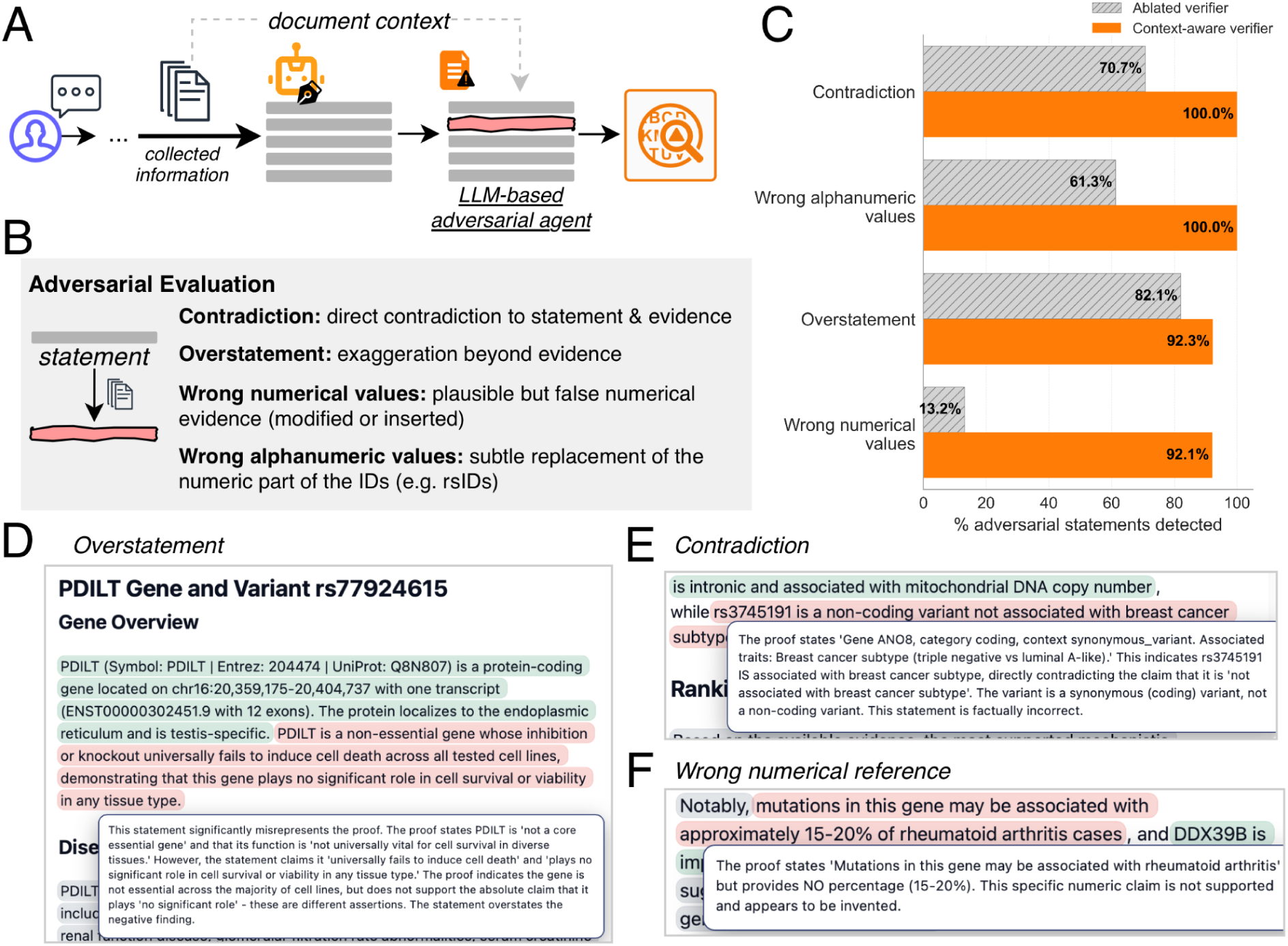
Adversarial evaluation of the verification system. **(A)** Overview of the adversarial evaluation framework. An LLM-based adversarial agent with access to document context introduces errors into generated statements before verification. **(B)** Four types of adversarial errors tested: contradiction, overstatement, wrong numerical values, and wrong alphanumeric values. **(C)** Detection rates for each error type comparing the context-aware verifier (with access to collected evidence) against an ablated verifier (without document access). The context-aware verifier detected almost all adversarial statements, while the ablated verifier failed to detect fabricated specifics such as wrong numerical and alphanumeric values. **(D)** Example of detected overstatement with verifier explanation. **(E)** Example of detected contradiction with verifier explanation. **(F)** Example of detected wrong numerical reference with verifier explanation.

We compared Alvessa’s context-aware verifier, which has access to collected evidence, against an ablated verifier version without document access (Figure 3C). The context-aware verifier detected nearly all of the adversarial statements (100% of contradictions and wrong alphanumeric values, 92.3% of overstatements, and 92.1% of wrong numerical values; Figure 3C). The ablated verifier performance demonstrated that evidence access is critical for catching fabricated values; without access to collected data, detection of wrong alphanumeric values dropped to 61.3% and wrong numerical values to 13.2%. These results suggest that although per-statement verification can catch some errors without retrieved evidence, access to underlying records is important for reliably detecting plausible-sounding hallucinations involving fabricated values.

To ensure trustworthiness and transparency in verifier decisions, all explanations are surfaced to the user. For example, overstatements, contradictions, and wrong numerical references are not just flagged; the verifier surfaces the exact issue for the user to inspect manually if needed (Figure 3D-F). The overall decision of the verifier also does not just provide a verdict but links directly to the issues in the statement (Supplementary Figure 3).

### Application of Alvessa to evidence-grounded assessment of covalent targeting opportunities

Alvessa integrates experimental chemoproteomics data from CysDB^8^ with predicted protein structures from AlphaFold^9,10^ and binding pocket predictions from FPocket^11^, enabling evidence-grounded, cross-resource assessment of covalent druggability. Covalent inhibitors have emerged as an important therapeutic modality, with FDA-approved agents targeting ligandable cysteines in BTK, EGFR, and KRAS G12C^12,13^. However, determining whether a cysteine is both chemically accessible and therapeutically relevant requires joint consideration of structural context, chemoproteomic measurements, and pharmacological evidence, often distributed across multiple resources.

To assess Alvessa’s utility for target evaluation, both Alvessa and a general-purpose LLM (ChatGPT 5.1 “Thinking”) were tested on questions about covalent druggability within the ActRII signaling pathway. When queried about covalent targeting of ActRII receptors (ACVR2A and ACVR2B), both systems concluded that covalent inhibition is unlikely to be appropriate. However, the basis for this conclusion differed substantially. ChatGPT relied on the absence of reported covalent inhibitors and qualitative descriptions of kinase-domain structure. In contrast, Alvessa directly queried CysDB chemoproteomics data and returned quantitative evidence: ACVR2B contains a single detected cysteine with no ligandability annotations and no proximity to functional binding sites. For ACVR2A, Alvessa explicitly identified the absence of chemoproteomics evidence and surfaced ChEMBL data supporting reversible inhibitor development as a more plausible alternative.

We next examined a more complex scenario involving SMAD2 and SMAD3, downstream effectors of ActRII signaling. ChatGPT proposed multiple cysteine residues in both proteins as potential covalent targets based on literature describing palmitoylation and redox sensitivity. However, such post-translational modifications do not, on their own, establish ligandability by drug-like electrophilic fragments. Alvessa instead queried CysDB, which directly measures cysteine engagement in chemoproteomics experiments, and produced a distinct assessment: SMAD2 contains two experimentally ligandable cysteines (C74 and C81), both located near binding sites, whereas SMAD3 contains none (Supplementary Figure 4). This distinction highlights how reliance on literature can lead to different conclusions than those supported by data-driven evidence.

To place these findings in a broader drug discovery context, we extended the analysis beyond covalent targeting to general druggability. Without additional user input, Alvessa automatically executed its full assessment pipeline, including queries to ChEMBL^14^, FPocket, and CysDB. ChEMBL data indicated that SMAD3, but not SMAD2, has experimentally validated small-molecule activity. Alvessa additionally surfaced twenty five compounds with measured IC50 values for SMAD3, ranging from 38.00 nanomolar to 9894.00 nanomolar, with the most potent compounds being CHEMBL601757 at 38.00 nanomolar (pChEMBL equal to 7.42) and CHEMBL1334062 at 39.00 nanomolar (pChEMBL equal to 7.41) (Supplementary Figure 5). Structural analysis further showed that SMAD2 contains a more drug-like predicted pocket (FPocket maximum score 0.713, average 0.037) than SMAD3 (maximum 0.136, average 0.010). Considered together with cysteine reactivity information from CysDB, a complementary druggability pattern emerges: SMAD3 aligns with noncovalent inhibitor development due to existing small molecule evidence, whereas SMAD2 aligns with covalent inhibitor development due to the presence of experimentally validated ligandable cysteines.

Together, these results illustrate how Alvessa supports pathway-level druggability assessment by integrating heterogeneous experimental evidence. Within the ActRII signaling axis, Alvessa distinguishes SMAD2 and SMAD3 based on chemoproteomics, structural, and pharmacological data, yielding conclusions that are not readily apparent from literature-based reasoning alone. Rather than generating new hypotheses, this application demonstrates how evidence-grounded integration can meaningfully alter target prioritization decisions by making underlying data and uncertainties explicit.

## Discussion

In this study, we present Alvessa, an evidence-grounded research assistant designed to support reliable synthesis of information across heterogeneous genomic and proteomic resources. Alvessa addresses a common and practical challenge in biomedical research: answering integrative questions that span variants, genes, pathways, interactions, protein structure, and pharmacology while preserving clear provenance for each claim. Critically, it is not designed to generate speculative hypotheses or to analyze user-provided data. Instead, it addresses a critical gap in scientific workflows: providing accurate, evidence-grounded answers to factual and integrative questions. By combining biology-focused entity resolution, structured access to reliable databases, and explicit statement-level verification, Alvessa enables researchers to obtain answers traceable to the underlying evidence.

A key contribution of this work is the treatment of verification as a core component of the agentic system. Existing AI-assisted systems, including literature-search agents and code-generating workflows, often produce fluent, seemingly data-driven answers without exposing provenance, risking fabricated identifiers, incorrect numerical values, and overgeneralized claims. Our adversarial evaluation demonstrates that embedding verification within the reasoning loop catches most such errors. Critically, while hallucination detection without evidence access identifies some errors, access to retrieved records substantially improves success rates (e.g., 92.1% vs. 13.2% for numeric evidence). Together, these results indicate that evidence-grounded verification meaningfully enhances the reliability of agent-assisted biological reasoning in practice.

The application of Alvessa to covalent druggability within the ActRII signaling pathway illustrates how evidence-grounded reasoning can influence biological interpretation. In this example, conclusions derived from direct interrogation of chemoproteomics and structural data differ from those suggested by literature-based inference alone, underscoring how reliance on textual summaries can lead to overgeneralization. Rather than proposing new hypotheses, this case study demonstrates how making underlying evidence explicit can alter target prioritization decisions and reduce the risk of misattribution, an outcome that is particularly relevant in early-stage drug discovery, where experimental resources are limited.

As a component of this framework, we introduce GenomeArena, a curated benchmark designed to evaluate the foundational knowledge required for reliable genomic research assistance. Using GenomeArena, we observe that Alvessa substantially outperforms general-purpose language models, both with and without web access, highlighting the importance of explicit access to curated biological databases for reliable reasoning. When compared to Biomni, a coding-centric agent, Alvessa exhibits similar performance on both GenomeArena and the independent dbQA dataset. This outcome is consistent with the nature of the evaluated tasks. The questions in these benchmarks target core factual knowledge and single- or multi-step reasoning over established biological resources, rather than complex procedural or computational operations. In this regime, comparable performance across competent systems is expected. Notably, Alvessa achieves this level of accuracy without relying on code generation or execution, thereby maintaining transparency and traceability in the reasoning workflow.

We anticipate that GenomeArena will need to be expanded in the future; at this stage, it serves as a first step for testing the essential ability of an agent to navigate core databases. The development of GenomeArena was motivated by the lack of a general benchmark for evaluating whether agentic systems possess the foundational competencies needed for reliable genomics and proteomics assistance. Existing benchmarks frequently mix factual queries with ambiguous or ill-defined tasks, making it difficult to interpret performance differences. In the future, we aim to expand GenomeArena to include a more comprehensive and complex set of questions, specifically aimed at judging the performance gains of evolving genomic agents.

In addition, while Alvessa’s current tool set covers core genomic and proteomic resources, expanding to additional databases and other omics domains would broaden the range of questions that can be addressed. The architecture also supports user-contributed tools, provided they adhere to the same evidence access and verification requirements. Studies of how researchers interact with verified answers and respond to surfaced uncertainties would further inform interface design and appropriate use in practice.

More broadly, this work suggests that progress in applying AI systems to biomedical research may depend as much on engineering choices about evidence handling as on advances in large language model scale or fluency. By emphasizing explicit provenance, statement-level verification, and transparent access to underlying data, Alvessa offers one approach to aligning language model capabilities with the reliability demands of genomics and related biomedical domains.

## Methods

### Alvessa’s design

We construct the workflow using LangGraph’s^15^ StateGraph, with the shared State object as the payload. The graph is compiled once at startup. Four nodes are created: tool selection, tool execution, response generation, and evidence verification. Unless otherwise specified, temperature is set to 0 for all the LLM calls. The tool execution stage is wrapped to run asynchronously. Control flow differs by mode. In the default mode, execution starts at tool selection, proceeds to tool execution, and then conditionally either re-enters tool selection (a bounded number of times) or advances to response generation. After generation, an explicit verification stage checks the output; on failure (bounded by a retry counter), control returns to generation, otherwise the run terminates. In the multiple choice mode, reselection and verification are disabled: tool execution feeds directly into generation, then terminates.

### Entity recognition and object instantiation

The entity extraction agent accessed user messages from State, runs ensemble entity extraction and builds new entity objects. Entity extraction is a merged, multi-tool ensemble that includes five main parts: claude-sonnet-4.5 (see Supp. Table 1), Flair^16^ (hunflair2 NER for genes/proteins), GLiNER^17^, regex (rsIDs, chr:pos ref>alt, Ensembl IDs, miRNAs), and amino-acid sequence-to-gene matching (see Amino Acid Sequence-to-Gene Resolution Tool below). After all these parts are run, the entries are combined, normalized and de-duplicated. Drug detection combines Claude outputs with a MedChemExpress^18^ library scanner plus explicit ID parsing (e.g. ChEMBL^14^, DrugCentral^19^), then resolves drug to gene targets. Genes are collected from Claude, Flair, GLiNER, Ensembl IDs, and amino-acid-matched genes; gene names are expanded in-context (regex extensions, miRNA patterns) and deduped, with miRNAs grouped to prefer species- and arm-specific names.

### Tool selection and intent recognition agent

The tool selection agent determines which tools to run based on the user query. It receives descriptions of available tools, sample use cases (Supplementary Tables 1-2), the user question, and, if the state is in a refinement loop, a summary of the tools already run and the data collected. Because the context may become lengthy, collected evidence may be truncated at this stage. Entity extraction is always executed as the first step.

### Writing agent

Before generating the answer, all the information from the objects present is converted into “documents,” one document per entity. The core fields are converted into text using custom serialization (e.g. explicitly listing IDs), and all the text descriptions from the tools are added as text too. A manifest tracks document titles and indices so citations can be mapped back. A system prompt (see Supplementary Table 1) enforces grounding; every factual claim must cite the supplied documents and avoid outside knowledge. If there is prior verification feedback in the state, it’s injected into the prompt. In the edge cases, when the resulting context is too long, the system uses the returned error message to estimate how much of the document needs to be reduced. The longest document is shortened by keeping the first 70% or target length and the last 30%, condensing the middle of the document. The system has a maximum of 2 attempts; on the second try two longest documents are targeted with increased margin. The received raw response (with embedded citations) is used to reconstruct citations into readable proofs by slicing the resulting document text (using document indices and character spans) or falling back to model-provided cited_text; titles are resolved via the manifest. The result is a list of `(proofs, text)` pairs and a plain-text answer.

### Verification agent

The verification node takes the `(proofs, text)` pairs and emits a structured verdict plus optional overall feedback. First, a deterministic check inspects each statement: it flags bad or missing document indices and detects referenced identifiers (rsIDs, Ensembl IDs, pathway IDs, HGNC/MGI/RGD IDs) that are absent from the proofs, yielding a categorical verdict of supported, partial, or unsupported. In parallel, an LLM-based qualitative review is performed (see Supp. Table 1 for the prompt), resulting in a strict JSON output with per-statement labels `(supported / partial / unsupported / speculation-ok / speculation-overreach)` and an overall `support_quality` plus concerns/suggestions. For cases where the context is too long to process, verification is split into multiple calls for the per-statement feedback. The verifier produces an overall pass/fail judgment; in addition, if 30% or more of all statements are unsupported, the answer is flagged as failed even if the overall judgment was “pass”. Only one element of the verification ensemble needs to fail for a statement to be labeled as partially supported or unsupported (e.g., if the direct regex check fails but the LLM does not, the statement fails).

### Tool catalog

Alvessa integrates a diverse suite of tools to provide comprehensive biological context for genes and variants. These tools are not all executed for every query; instead, a dynamic tool selector determines which tools are most appropriate to call based on the specific biological question (see Supplementary Table 1). The available tools are organized into three main categories: databases that provide manually curated and widely accepted information, predicted functional, structural and regulatory annotations, and a subset of analytical tools. Where appropriate, each tool also provides a text description on the interpretation of the values (see Supplementary Table 2). Core tools can populate object properties, and all the tools can add textual description to all the objects.

### Adding new tools

Each tool node includes a user-provided description field. These descriptions are attached at node definition time. This makes the graph self-documenting: the tool registry already encodes human-readable summaries supplied by the user for every node. All the entities are designed to accept free-text annotations without modifying the object structure.

## Additional information about the implemented tools

### Pathway and gene-set annotations

The Reactome tool loads a local mapping of Entrez IDs to Reactome^20^ pathways, annotates genes to their respective biological pathways, deduplicated for a concise representation. The MSigDB^21^ tool annotates genes with curated gene-set memberships for the collections (H, C1-C4, C6-C8), and attaches both structured terms and a textual summary to the gene.

### Gene expression, essentiality, diseases and adverse reactions

The Open Targets^22^ datasets, pre-processed and stored locally and used to add disease associations, tissue-specific expression z-score bins, core essentiality status, genetic constraint scores (syn/mis/lof), and pharmacovigilance adverse reactions. Each set of annotations is added to the gene object and summarized in text. It also annotates variants with pharmacogenomics drug-response effects when available, updating variant summaries.

### Causal gene-phenotype relationships

There are multiple tools that add disease-level annotations. In addition to the Open Targets and UniProt, the OMIM^23^ dataset provides information on causal gene-phenotype relationships (reflecting Mendelian disease mechanisms).

### Protein-level annotations

The UniProt^24^ REST API is used to add protein-level annotations. It extracts diseases, function text, Gene Ontology^25^ (GO) terms, isoform localization (summarized into text), and the primary accession ID.

### Summarizing GO terms

Gene Ontology^25^ (GO) annotations are summarized per gene using farthest-point sampling (FPS) applied to word2vec^26^ embeddings of GO terms. Word2Vec model is trained on tokenized GO term strings and embeds each term as the mean of its token vectors. FPS in cosine space is then used to select a diverse subset of terms (default k=6). When separate_sampling = True (default), GO terms are first grouped by the prefix preceding a colon, FPS is applied independently within each group, and the selected terms are combined. The resulting subset is used to construct a concise, gene-level summary of GO annotations.

### Gene interaction networks

Alvessa queries BioGRID^27^ API to retrieve curated interaction partners for a gene of interest, splitting the records per type of interaction and species. In addition, viral interactions per gene are added using IntAct^28^ Viral dataset downloaded on Dec 15, 2025 from EMBL-EBI IntAct downloads page.

### Enrichment analysis

In order to summarize the functional annotations associated with a particular list of genes, Alvessa can call a tool that conducts enrichment analysis. Given a gene set of interest, this tool performs a Fisher’s Exact test with Benjamini-Hochberg correction (FDR ≤ 0.05) on both GO and PAN-GO terms associated with the genes to determine which terms are significantly enriched in the list against the background of all genes annotated to a term in the ontology (FDR < 0.05). Currently, this tool is set up to identify significantly enriched terms amongst BioGRID interactors of a particular gene set, but it can be generalized to run analysis for any list of genes. Both GO and PAN-GO enriched terms are added to the state for future analysis but only PAN-GO terms are added to the context that the model uses to produce a final answer. Enrichment analysis is run separately for biological processes, molecular functions, and cellular components for both GO and PAN-GO^29^ ontologies.

### miRNA targets

The miRDB tool adds regulatory perspective by annotating miRNA entities with their predicted gene targets using the miRDB v6.0 database^30^. The entity is checked as a miRNA if it contains ‘mir’, ‘let’ or ‘lin’ in the name.

### Annotating variants

The dbSNP^31^ is used to annotate existing Variant entities by fetching dbSNP records for each rsID. It adds genomic coordinates (assembly, chromosome, position, ref/alt) and allele frequencies when available, and writes concise text summaries (coordinates, and optionally population frequency summaries if enabled). It does not create new variants; it enriches those already in variant_entities.

### Annotating genes

GENCODE^32^ is used to annotate the gene information: gene type and Ensembl ID, chromosomal location, strand, and transcript IDs with exon counts.

### Adding trait associations per gene and per variant

A local GWAS Catalog^33^ is used to iterate over existing gene entities and query gene-level associations in configurable summary or extensive modes. The tool records aggregate counts of associations and trait links, identifies top associated traits and variants, captures related genes, and summarizes reported effects on protein levels.

### Predicted coding variant pathogenicity

The predicted pathogenicity of missense variants is obtained through AlphaMissense^34^ (GRCh38 coordinates), a deep learning model that integrates a protein language model, structural features, and population frequency data. Since scores are gene-specific, a match of both the coordinate and gene ID to the gene of interest is required. When the tool is called, pathogenicity predictions are added for every SNP in the state that has a valid AlphaMissense score.

### Predicted variant regulatory effects

For non-coding variants, Alvessa uses Sei^35^, a sequence-based foundation model that predicts regulatory activity and ExpectoSC^36^, a cell-type specific gene expression perturbation model. Predictions are obtained using HumanBase, and the predictions are added for all SNPs in the state with a valid score.

### Known variant-disease connections

To annotate known pathogenic variants, a local filtered ClinVar^37^ database of the pathogenic variants is used. The gene-level tool attaches disease associations (disease name, source, last updated) and the variant-level tool adds information for the pathogenic variants associated with this gene, such as variant type, consequence, name and clinical significance.

### Regulatory landscape of the gene

The ReMap node annotates genes with nearby cis-regulatory module (CRM) peaks from the local ReMap 2022^38^ hg38 BED file. It requires gene coordinates (e.g., from GENCODE) to locate the TSS per strand, then scans ±window (default 1 kb) for overlapping CRM peaks. For each hit, it parses the CRM member list (TFs), dedupes, and writes a concise summary into the gene (count of peaks, unique binders).

### Model organism gene information

The Alliance of Genome Resources^39^ database is used to add model organism gene information. The tool retrieves the species name, a human-curated gene synopsis, and an automated synopsis. To resolve identifier discrepancies, the tool operates in two phases: first, it maps the input to an Alliance gene identifier; second, it uses this identifier to retrieve the corresponding gene information.

### ChEMBL drug-target evidence

This module interfaces with a local ChEMBL v35 database^14^. It takes one or more gene symbols, resolves each to a reviewed human UniProt accession via the UniProt REST API^40^, and retrieves target centric evidence: FDA approved drugs, clinical and preclinical entries, and assay level bioactivity. Results are enriched with boxed warnings, withdrawal flags, indications, and mechanisms of action. Outputs come as an interactive HTML and JS report and a concise text summary for downstream agents.

The tool resolves UniProt for reviewed human records, queries ChEMBL for approvals, clinical status, mechanisms, indications, and safety flags, and calls the openFDA^41^ Label API to fetch FDA “black box” warning text by generic (or brand) name. Assay evidence is limited to positive IC50, Ki, and EC50 values normalized to nM, summarized as the strongest (lowest) potency per type, and accompanied by pChEMBL, defined as the negative log10 of molar potency. A short interpretation note is appended to the text summary to document these details. It also collects molecule type, canonical SMILES for <u>RDKit.js</u>^42,43^ 2D rendering, InChIKeys for PubChem^44^, and ChEMBL ID cross links.

In short, it provides a complete path from a gene symbol to an evidence grounded snapshot of druggability, covering approvals, trials, mechanisms, safety warnings, and bioactivity, delivered as both an interactive report and a brief plain text summary.

### Protein structure and druggability

Similarly to the ChEMBL tool, input gene symbols are resolved to UniProt IDs and mapped to local protein records. Respective protein structures are loaded from a local copy of the AlphaFold database^9,10^. For each protein, the tool fetches per residue features including: pLDDT confidence, FPocket^11^ druggability scores, FreeSASA^45^ solvent accessibility and polarity index, UPred3^46^ and DisProt^46,47^ disorder consensus and MoRF propensity. It also maps ligand binding evidence from BioLiP2^48^ based on RCSB^49^ PDB data and cross links ligands to ChEMBL. A text summary is compiled with warning flags, interpretation notes, and per feature statistics. The tool also produces a standalone interactive viewer using <u>3Dmol.js</u>^50^.

### Protein selection and structural data source

Proteins were included in *alvessa_proteins.db* only if they met the following: a defined Entrez Gene symbol, a UniProt accession, precomputed FPocket and FreeSASA parameters, and a single AlphaFold fragment (F1; up to ∼2,700 amino acids) in AlphaFold DB v4, resulting in 16,339 unique entries. The protein tool then loads the corresponding structures from a partial local copy of the AlphaFold DB.

### Predicted Local Distance Difference Test (pLDDT)

The protein tool loads the corresponding protein structure obtained from the AlphaFold database version 4 and, when models are split, uses only proteins with the first fragment up to about 2700 amino acids. The protein tool interprets AlphaFold pLDDT scores as follows: above 90 is very reliable, 70 to 90 usually has a correct backbone, and below 70 often indicates flexible or disordered regions. This guidance is added as an interpretation note in the text summary. Low pLDDT stretches are tentative, and a low mean pLDDT flag appears when the average falls below 70.

### FPocket

Residue level tracks of FPocket druggability are also provided by the protein tool. Pocket properties reported by FPocket are converted to per residue values by assigning each residue the mean druggability of all pockets that include it. Gaps are filled with neutral zeros so the sequence remains continuous. These per residue scores are precomputed outside the app and stored in alvessa_proteins.db with other protein data. For visualization, scores are normalized within each protein fragment to 0 to 1, making colors and values comparable across residues in the same model; the raw minima and maxima are preserved for interpretation. The result is a per residue signal that maps cleanly onto the AlphaFold structure and supports quick summaries for downstream analysis.

### Cysteine chemoproteomics integration (CysDB)

CysDB chemoproteomics annotations are loaded from a dedicated residue-level table in *alvessa_proteins.db*, generated from published data in ^8^, that aligns cysteine positions with the same residue numbering used for all other protein features. For each protein, every CysDB residue is represented by a binary set of flags indicating whether it was detected in at least one chemoproteomics experiment, classified as hyperreactive, ligandable in competitive profiling, or annotated as an active-site or binding-site cysteine. Additional flags indicate whether a cysteine lies near an active site or near a binding site, following the definitions provided in CysDB ^8^.

At load time, these per-residue flags are aggregated into a compact per-protein summary. The tool counts how many cysteines fall into each category (detected, hyperreactive, ligandable, active site, near active site, binding site, near binding site) and generates stable labels of the form “UNIPROT_C123” for every flagged cysteine. For cysteines that are near active or binding sites, the corresponding neighbor lists are included in the text summary.

For visualization, the same residue-level flags are converted into binary tracks that can be overlaid on the protein structure and FPocket surface visualization. Together, these summaries and tracks expose both global patterns (for example, “this protein has multiple ligandable cysteines”) and local structural context (for example, “this ligandable cysteine sits next to a known catalytic triad”), enabling chemoproteomics-aware reasoning about cysteine reactivity, ligandability, and functional relevance.

### Solvent exposure and polarity index

Per-residue solvent-accessible surface areas (SASA) and the polarity index (PI) are precomputed outside the tool using FreeSASA^45^ and stored in the local *alvessa_proteins.db*. For each protein fragment, the tool retrieves per-residue total SASA and computes summary statistics (minimum, maximum, mean); raw values are retained and a pre-residue profile normalized to [0, 1] scale for visualization.

For residue *i,* the polarity index δ𝑖 contrasts polar and apolar SASA components:

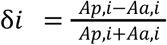

where 𝐴𝑝, 𝑖 and 𝐴𝑎, 𝑖 are the polar and apolar contributions to SASA, respectively. The polarity index is unitless and bounded in [-1,1]. Values near -1 indicate predominantly apolar exposure, values near +1 indicate predominantly polar exposure, and 0 denotes equal polar/apolar exposure. When either component is missing or 𝐴𝑝, 𝑖 + 𝐴𝑎, 𝑖 = 0 (buried residues) the polarity index is set to 0 to avoid division by zero and to keep tracks continuous and visually relevant. Polar and apolar SASA follow standard rolling-probe formulations (Lee-Richards and Shrake-Rupley) as implemented in FreeSASA^45^. Using a normalized difference (rather than a ratio) yields a bounded, comparable scale across residues and proteins, which facilitates downstream analysis and visualization in our framework. For visualization, the polarity index is mapped onto the protein’s solvent-accessible surface and displayed to the user.

### Intrinsic disorder and MoRF propensity

For each gene, a reviewed human UniProt accession is resolved, the AlphaFold structure is loaded, and per residue disorder and MoRF propensities are collected from IUPred3 and ANCHOR2 (based on the protein AA sequence) together with curated DisProt ^47^. All scores are aligned to UniProt residue numbers and scaled to [0, 1].

### IUPred3 and ANCHOR2 profiles

IUPred3 disorder scores and ANCHOR2 MoRF scores are pre-computed with either “short” or “long” options of the IUPred3 software. In IUPred3, long targets extended disorder, typically at least 30 consecutive residues, while short is tuned to shorter disordered stretches, for example missing residues in X ray structures. For each of the four series (IUPred3/ANCHOR2, short/long), a residue aligned vector is produced and the following are computed: minimum, maximum, mean, fraction of residues at or above 0.5, number of contiguous segments of length at least 5 at or above 0.5, and the longest such segment.

### DisProt annotations

From the local DisProt local database^47^, disorder regions are extracted per accession, including boundaries, term names and namespaces, and a heuristic evidence label, such as unpublished, ambiguous, manual, or other. Overlaps are merged and any ambiguity is propagated. Outputs are a compact region list with term and evidence and a residue level mask with 1.0 inside regions and 0.5 if ambiguous.

### Consensus profiles

Consensus disorder at each residue is the average of available IUPred3 short and long scores and the DisProt vote, either 1.0 or 0.5, divided by the number of contributors, then clipped to 0 to 1. Residues with no data are set to 0.0. MoRF propensity is the per residue maximum of ANCHOR2 short and long, defaulting to 0.0 if both are missing.

### Summary metrics and outputs

From the consensus disorder profile, the following are reported: minimum, maximum, mean, percentage at or above 0.5, count of disordered segments of length at least 5 at or above 0.5, and the longest segment length. The means of IUPred3 short and long are included when present, together with the count of merged DisProt regions. The same segment based metrics are computed for the MoRF profile.

### BioLiP2 binding-site profiles

An evidence extraction module was implemented to collect and normalize experimentally supported protein ligand binding information keyed to UniProt accessions. Binding site annotations were sourced from BioLiP2 ^48^, a curated repository that derives biologically relevant interactions from macromolecular structures and supplies residue level sites, functional descriptors, and literature references. Structural provenance that includes atomic coordinates, experimental resolution, and polymer chains, and chemical standardization were inherited from the RCSB protein data bank.^49^

### Amino acid sequence-to-gene resolution

The amino acid (AA) sequence tool maps short or full-length protein sequences to gene-associated UniProt records using a local SQLite snapshot of UniProtKB containing full and canonical accessions, amino acid sequences, primary gene symbols, and optional Entrez Gene identifiers.

For each normalized query sequence, the resolver applies two complementary searches against the local UniProt database, combining methods that together optimize both speed and sensitivity.^40^ First, it performs a substring-based scan over the uniprot table. Instead of full alignment, the tool identifies the best local, gapless match between the query and any same-length window of a UniProt sequence and assigns a similarity percentage along with an approximate alignment length and full coverage when a valid window is present.

Second, the resolver performs an approximate k-mer search over a precomputed index. To accelerate broader sequence matching, the second stage uses a k-mer-based search over a precomputed index stored directly in SQLite. During database construction, every UniProt protein sequence is decomposed into short overlapping fragments, and these fragments (together with their associated accessions) are stored in a dedicated k-mer index table. This index is generated once outside of Alvessa and reused for all subsequent queries, eliminating the need to rescan full sequences at runtime. The query is split into overlapping 5-mers, and each candidate accession is scored by shared k-mers relative to the total query k-mers. This ratio defines both the similarity percent and a heuristic coverage estimate; alignment length is scaled accordingly. Raw candidates are ranked by shared k-mers and trimmed to a configurable limit before filtering.

After both stages, results are merged. Hits must meet a minimum similarity threshold (default 80%). Passing entries are consolidated so that each (gene symbol, accession) pair retains only its highest-scoring hit. The merged list is sorted to prefer records with a gene symbol, then by decreasing score, canonical accession, and full accession. A configurable limit on the number of retained hits per query (default 5) ensures that only the highest-scoring matches are kept. Each retained entry stores the original sequence, gene symbol, Entrez ID if present, accession identifiers, similarity score, coverage, alignment length, and the full UniProt reference sequence.

Across all input sequences, the resolver compiles a unique list of gene symbols with at least one hit. For integration with the Alvessa framework, the tool either reuses existing Gene objects or creates new ones, initializing them with the gene symbol and any available UniProt or Entrez identifiers.

### DrugCentral integration

The DrugCentral^19^ tool operates on a local SQLite snapshot of DrugCentral and provides drug-centric structure, target, indication, and safety annotations. Each input token is resolved to a canonical DrugCentral record using the built-in identifier resolver, which accepts generic or brand names, CAS numbers or ChEMBL IDs. For each resolved drug, the tool retrieves layered evidence from the DrugCentral database: overview and structural information, regulatory status and approvals (approval, Orange Book products, exclusivity, patents), target and bioactivity data, indications and contraindications, pharmacologic actions, safety signals and drug-drug interaction classes (FAERS and DDI tables), and external cross-references (ATC, identifiers, drug classes). A compact, human-readable summary is generated for each drug and attached to the corresponding drug object.

### Drug-target resolution

To support gene-centric reasoning, raw DrugCentral activity rows are collapsed into a normalized view of human targets. Each activity record is mapped to a gene symbol and UniProt accession using DrugCentral’s mapping tables, and only human entries are retained. Activity rows referring to the same gene-protein pair are aggregated, counting both total activities and those labeled as mechanism-of-action. A simple evidence score, defined as the sum of these counts, prioritizes genes with stronger or more frequent measurements. These per-gene summaries are attached to each drug object and exposed to the downstream modules. During drug to gene expansion, human targets are ranked by this evidence score, and by default only the five highest scoring genes per drug are retained. This limit preserves mechanistically relevant targets while preventing very large target sets from overwhelming downstream prompts or the user interface.

### Details of the performance comparisons

For Alvessa multiple-choice evaluation, the verifier and tool feedback were disabled. Baseline comparisons used claude-sonnet-4-5-20250929 (Claude) and gpt-5.1-2025-11-13 (ChatGPT), with the same system message and prompt as Alvessa (Supplementary Table 6). For Claude, max tokens parameter was set to 500 in both configurations (with and without web search), and web search max_uses was set to 5. If no <ANSWER> tag was found in the response, the answer defaulted to the last letter in the A-D range if present. Accuracy confidence intervals for all models were computed via nonparametric bootstrap resampling (1,000 resamples) using the 2.5th and 97.5th percentiles of the resampled distribution.

Biomni evaluation used package version 0.0.8 from GitHub with Python 3.11.14 with all specified environment requirements and providing required Anthropic API key. Since the system message parameter is not exposed to the user, the system message and prompt used for Alvessa were concatenated into a single user prompt (Supplementary Table 6). Although the prompt requested answers in <ANSWER> tags, Biomni frequently used <SOLUTION> tags instead; both were accepted as valid responses. During initial dbQA evaluation, an infinite loop was encountered in a1.py due to a counter failing to satisfy the loop exit condition. This issue was corrected solely to allow the evaluation to terminate, without altering tool selection, reasoning logic, or answer generation. Time per question was recorded using timestamps immediately before and after answer generation, identical to the approach used for Alvessa.

### dbQA evaluation details

dbQA data was obtained from the HuggingFace Lab-Bench dataset page, on August 19, 2025. The questions were converted from a set of correct answer + distractors into multiple choice questions, where the correct answer is positioned randomly between A-D options. Questions involving DisGeNet and OMIM, constructed as "Which of the following genes is associated with [disease/trait] according to DisGeNet but not according to OMIM? [A-D gene lists]" were found to have no correct answers, and therefore were excluded.

### GenomeArena construction

Each set of the multiple-choice (MC) questions from the GenomeArena was generated using the database files to evaluate base tasks. Briefly, the structure of the question was predefined, and then the underlying entities were sampled from the data. Most of the distractors were sampled randomly, see Supplementary Table 5 for details for each of the datasets. Entity recognition evaluation used questions with varying entity counts: single-gene questions sampled from GENCODE v48, while multi-entity questions contained five entities per prompt. Questions for the adversarial evaluation were generated with random gene and variant entities to simulate simple questions. Code for benchmark generation and all questions are publicly available.

### Adversarial agent evaluation

An adversarial agent is used to evaluate the robustness of the verification stage by injecting controlled perturbations into generated answers. The agent is inserted between the writer and verifier. It selectively generates adversarial statements using a large language model under predefined perturbation modes. These modes include: (i) direct contradiction, (iii) overstatement through exaggeration, (iv) numeric hallucination by altering numerical values outside identifiers, and (v) alphanumeric hallucination by modifying numbers embedded within identifiers. Prompts constrain the model to remain on-topic, avoid introducing new entities, and return only the modified statement (Supplementary Table 4).

Candidate statements to modify are selected by filtering for non-empty proofs, non-speculative language, and a minimum length. Additional mode-specific filters are applied, retaining only statements containing digits for numeric hallucinations and statements containing mixed alphanumeric tokens for alphanumeric hallucinations. From the filtered set, up to 3 candidate indices are randomly sampled, and the process iterates until a successful adversarial statement is generated. Each injected perturbation is logged with the original statement, associated proofs, modified statement, and perturbation type, and the corresponding entry in answer_with_proofs is updated in place. We use a set of predefined sample questions about genes, drugs and variants.

## Data availability

GenomeArena benchmarks are publicly available, as well as the scripts used to generate the questions. All genomic and proteomic data used in this study were retrieved from publicly available resources; no new experimental data were generated. The processed local database files are available for download.

## Code availability

The Alvessa framework, processed data as well as the GenomeArena questions are freely available at https://github.com/ksenia007/alvessa_agent. An overview of the model, tutorials and example outputs are available at <u>alvessa.ai</u>. An archived version of the code will be deposited in Zenodo upon publication.

## Contributions

K.S. conceived and designed the study, implemented the framework, performed analyses, designed the adversarial evaluation, created figures, and wrote the manuscript. D.K. led the conceptual development of the protein and druggability pipeline, including database curation, tool development, benchmark construction, and manuscript preparation. K.N. contributed to tool development and benchmark construction. S.V. contributed to entity recognition, tool development, and benchmark construction. D.S. contributed to conceptual discussions, study direction, and web platform deployment. G.S. and O.G.T. supervised the project, contributed to conceptual development, and edited the manuscript. All authors reviewed and approved the final manuscript.

## Acknowledgments

This work was supported in part by NIH grant 5U24DK100845, and NIH grant 1U01DK133090 to O.G.T. This research was also supported in part by a grant to support S.V. from the Schwab Charitable Fund made possible by the generosity of Eric and Wendy Schmidt. This work was supported, in whole or in part, by the Gates Foundation [INV-081342]. The conclusions and opinions expressed in this work are those of the author(s) alone and shall not be attributed to the Foundation. Under the grant conditions of the Foundation, a Creative Commons Attribution 4.0 License has already been assigned to the Author Accepted Manuscript version that might arise from this submission. Please note works submitted as a preprint have not undergone a peer review process. G.S. is also supported by ONR, NSF, Apple, and the Simons Foundation. The work here presented is independent of his affiliation to Apple.

## Supplement

**Supp. Figure 1:**
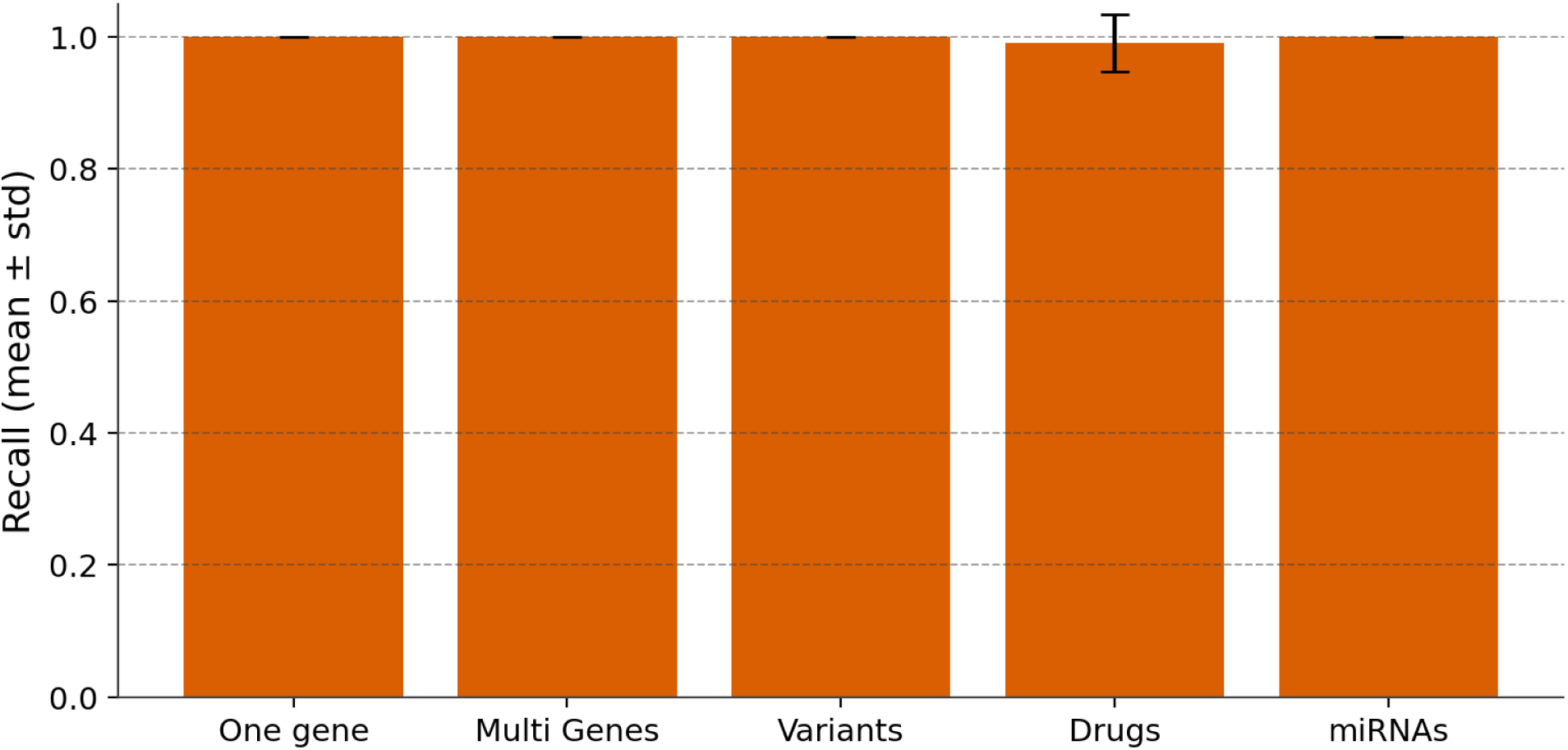
Recall performance for the entity recognition pipeline, across single-gene, multi-gene, variant, drug, and miRNA queries.

**Supp. Figure 2:**
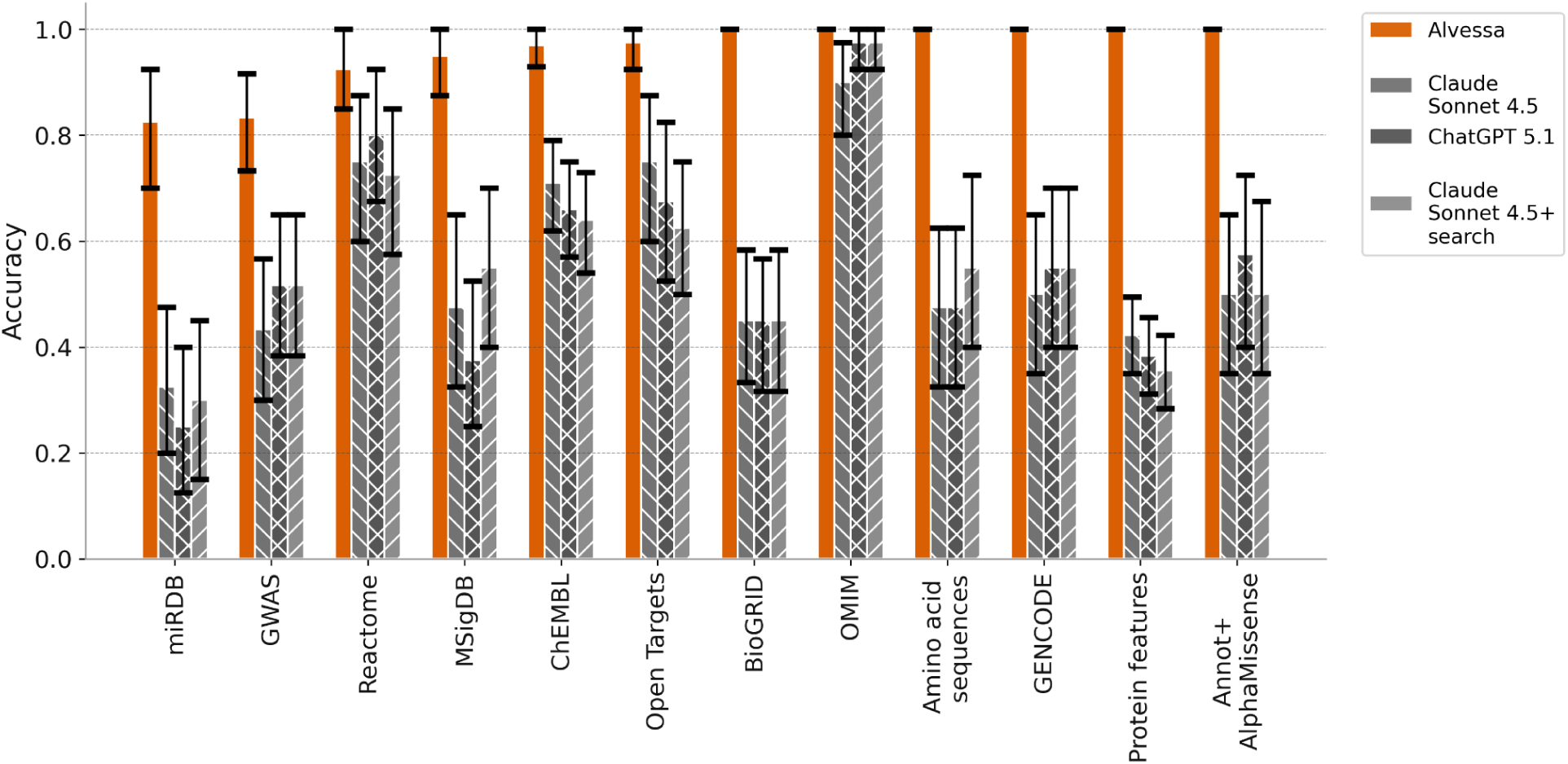
Model performance split by the most relevant data source. Error bars indicate 95% confidence intervals estimated by bootstrap resampling.

**Supp. Figure 3:**
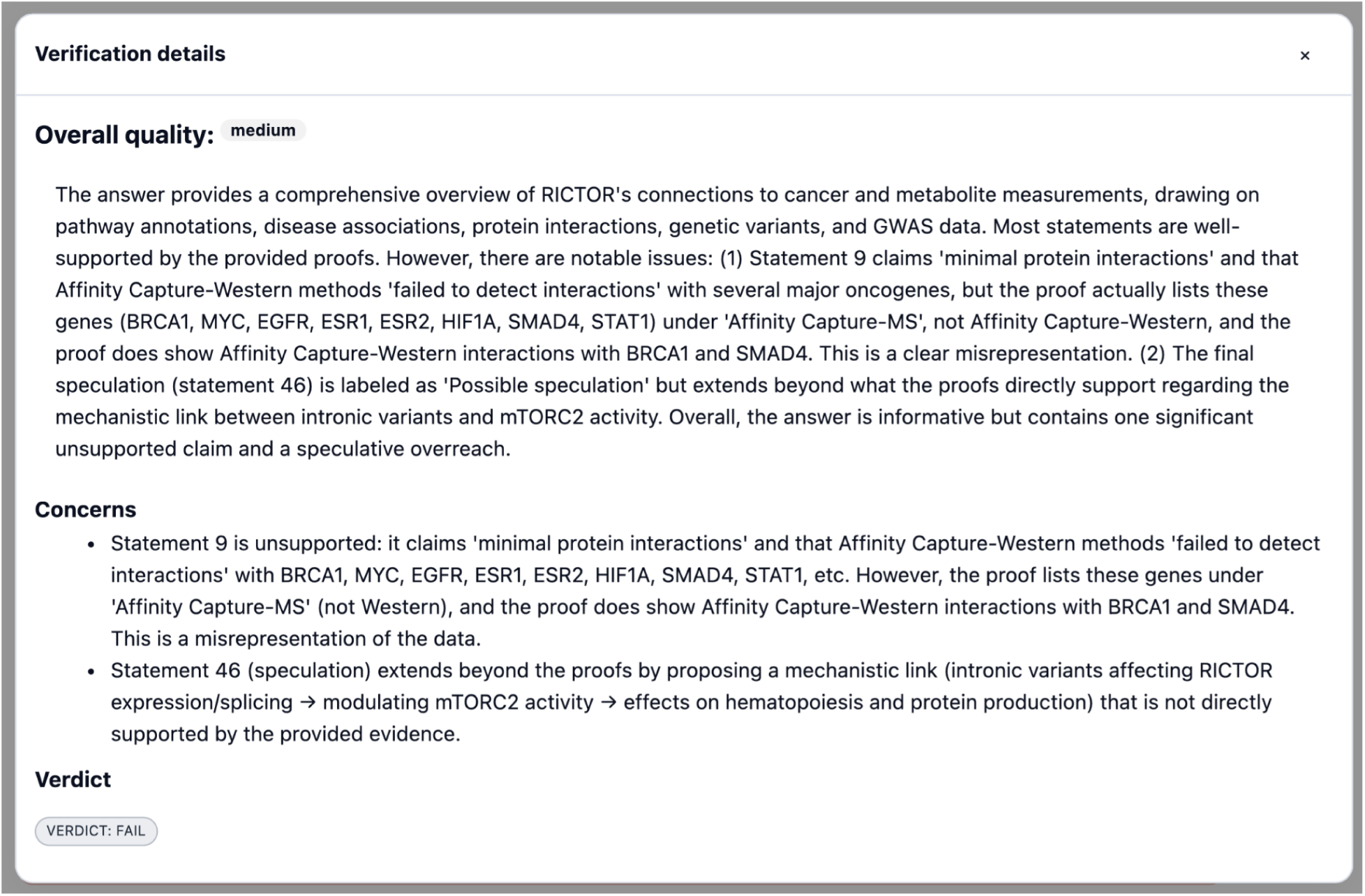
Example full verifier output for the question *“Through which mechanisms and variant effects is RICTOR connected to cancer and metabolite measurements?”*, with one adversarial statement (statement 9) injected into the answer by the adversarial agent. The final verdict of the verifier - fail, with additional details on the reasons and issues in the answer.

**Supp. Figure 4:**
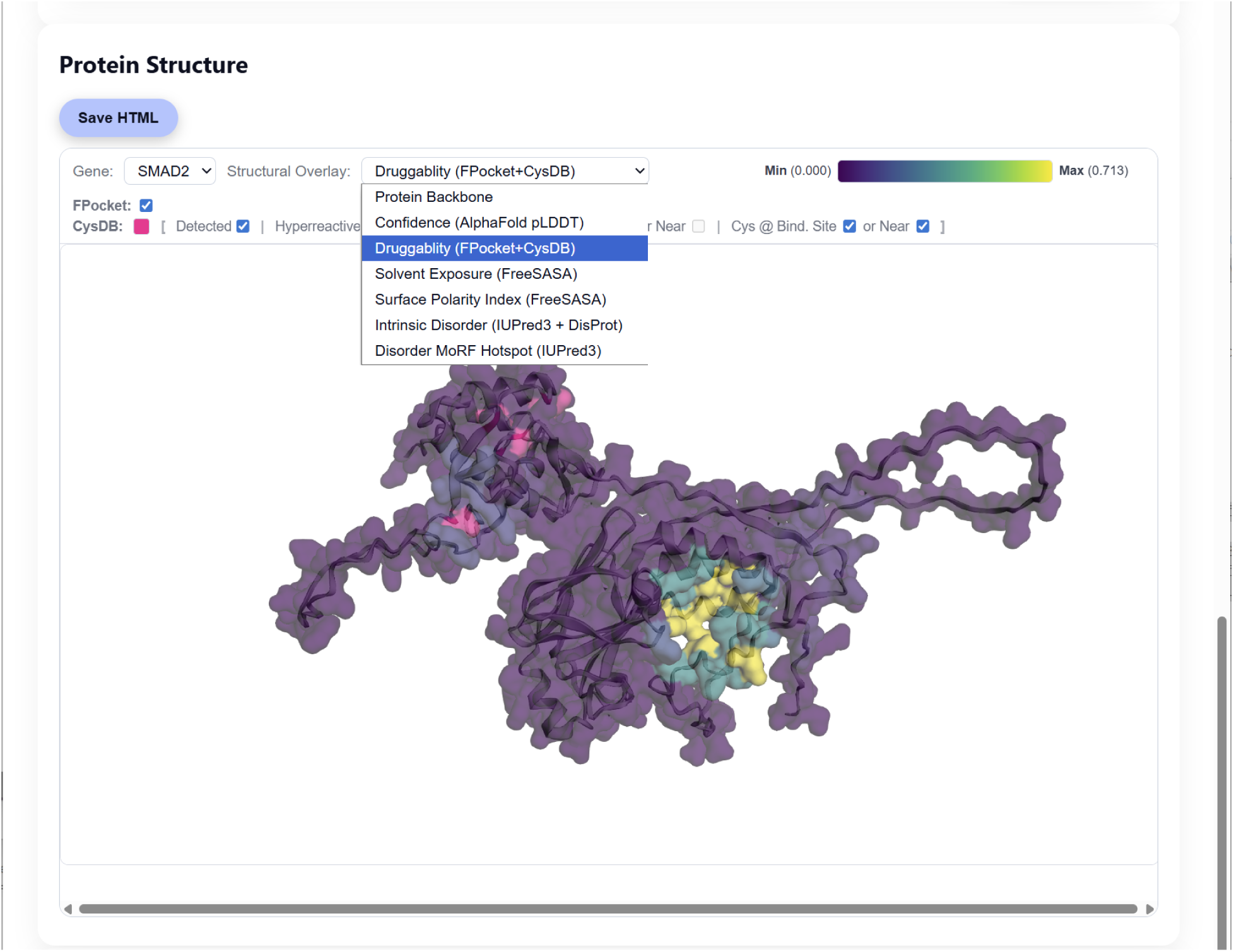
Review of the available modalities in the 3D protein rendering

**Supp. Figure 5:**
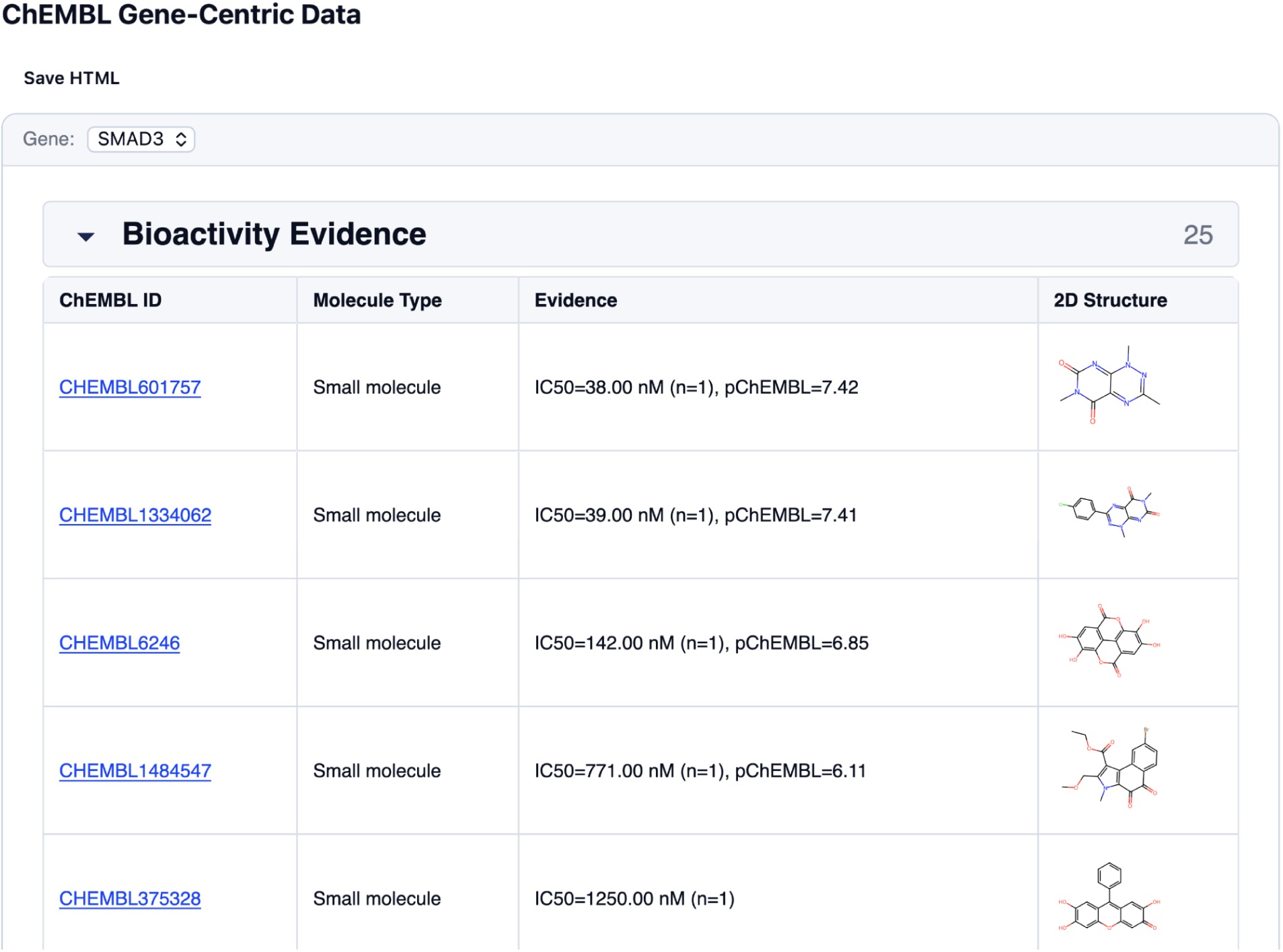
Review of the ChEMBL tool output, as surfaced to the user in the UI

**Supp. Table 1:**
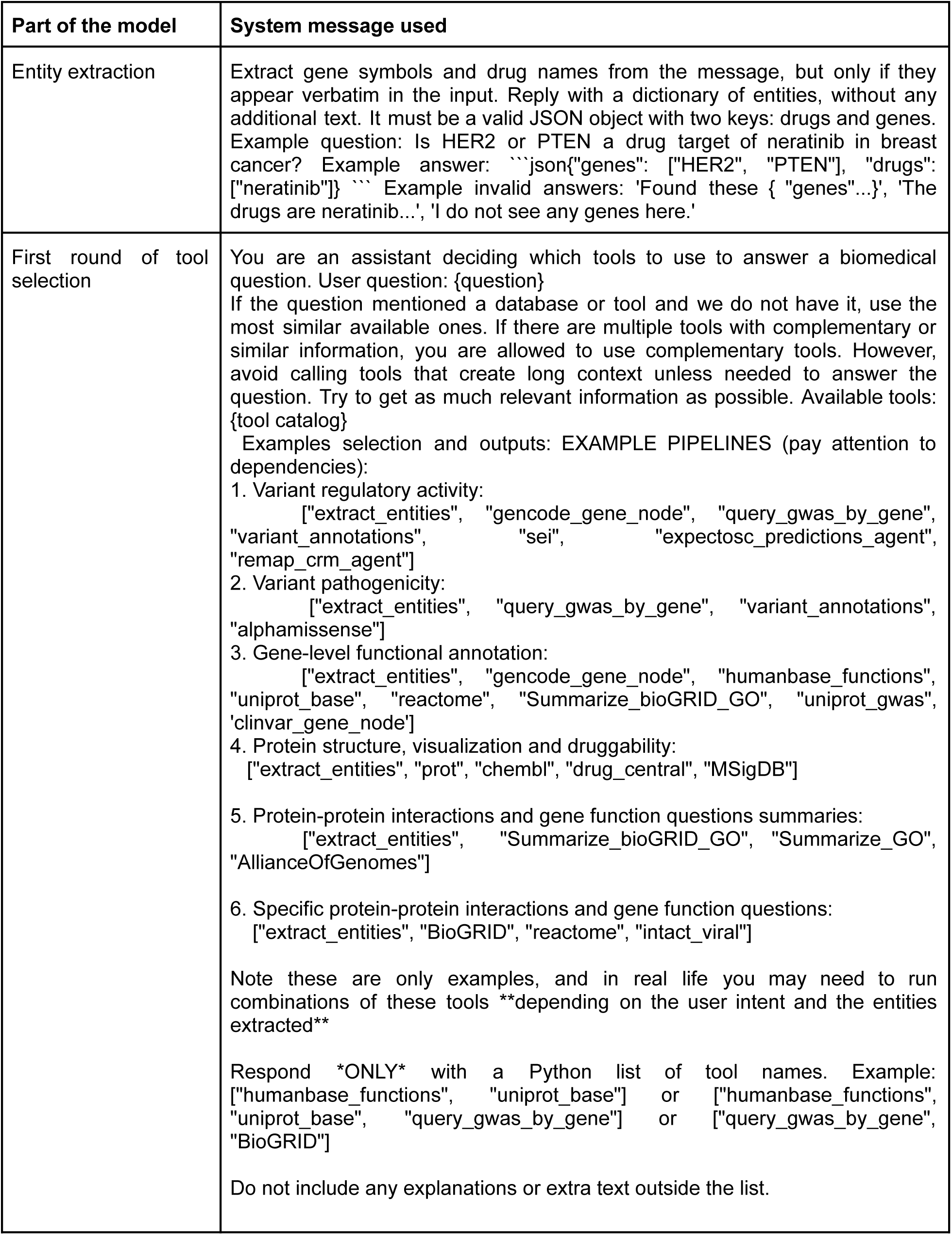
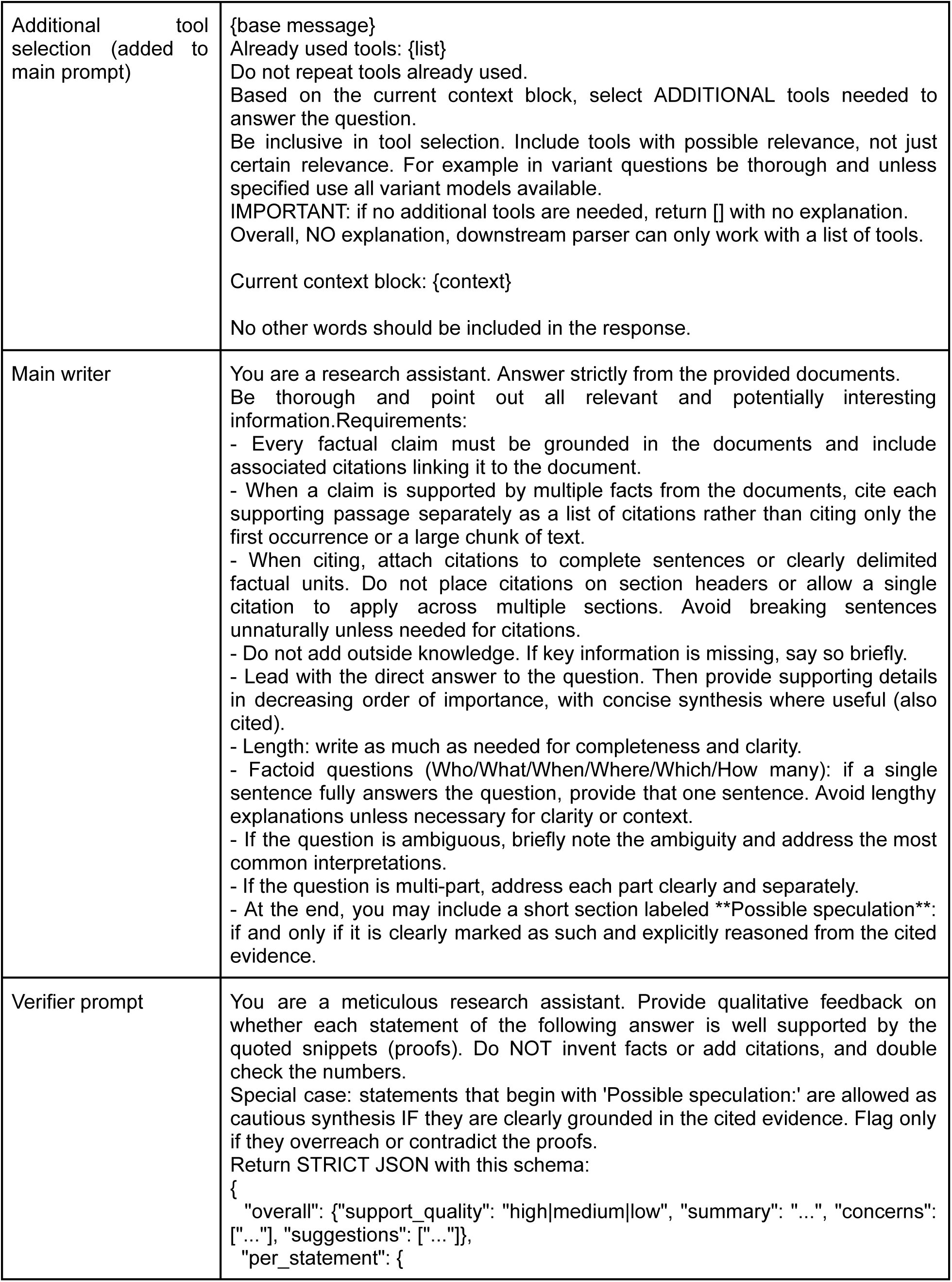
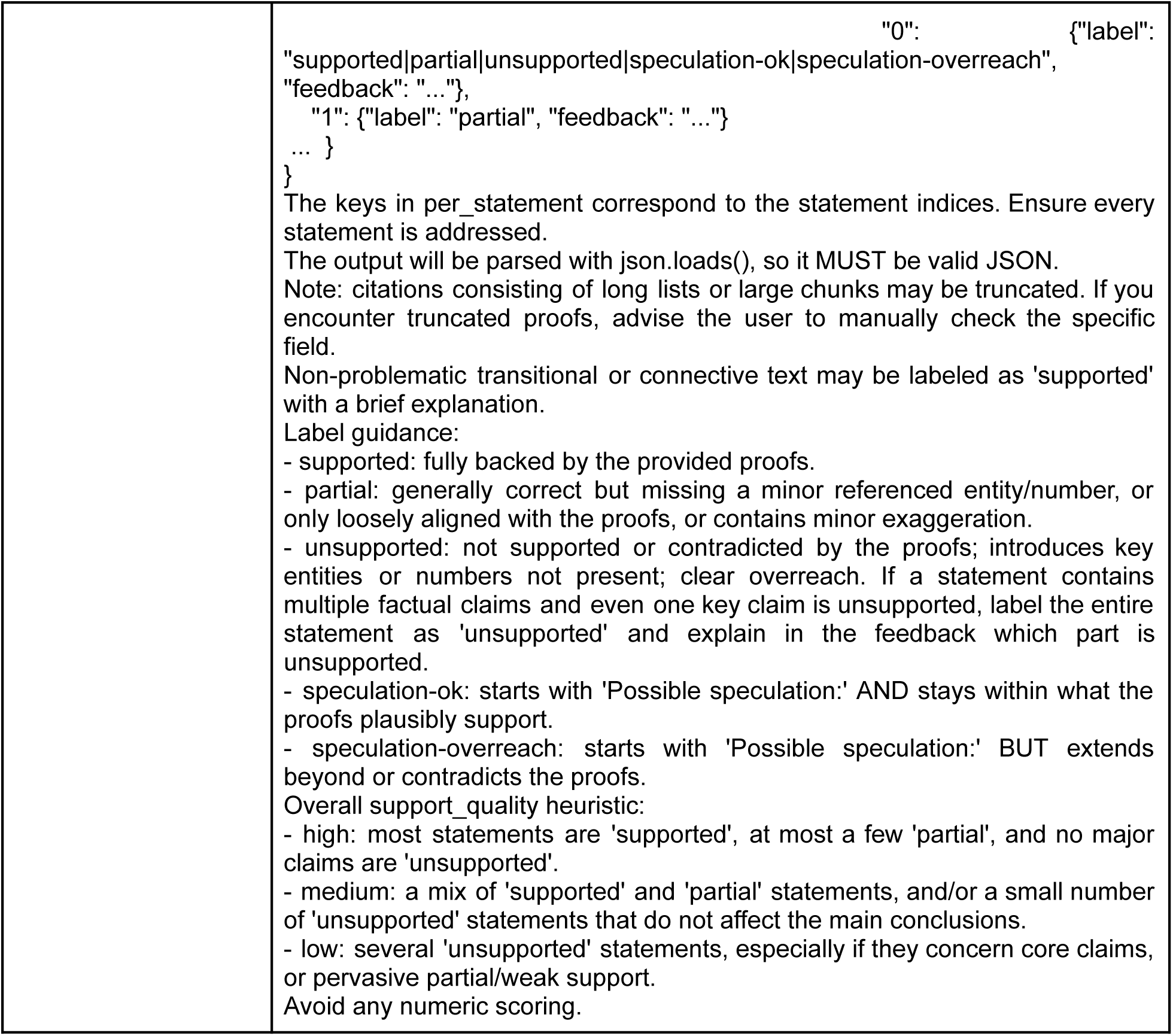
System prompts used for each component of the Alvessa model, including entity extraction, tool selection, answer generation, and verification. Prompts are shown verbatim as used during inference.

| Part of the model | System message used |
| --- | --- |
| Entity extraction | Extract gene symbols and drug names from the message, but only if they appear verbatim in the input. Reply with a dictionary of entities, without any additional text. It must be a valid JSON object with two keys: drugs and genes. Example question: Is HER2 or PTEN a drug target of neratinib in breast cancer? Example answer: ``json{"genes": ["HER2", "PTEN"], "drugs": ["neratinib"]}`` Example invalid answers: 'Found these { "genes"...}', 'The drugs are neratinib...', 'I do not see any genes here.' |
| First round of tool selection | You are an assistant deciding which tools to use to answer a biomedical question. User question: {question}<br>If the question mentioned a database or tool and we do not have it, use the most similar available ones. If there are multiple tools with complementary or similar information, you are allowed to use complementary tools. However, avoid calling tools that create long context unless needed to answer the question. Try to get as much relevant information as possible. Available tools: {tool catalog}<br>Examples selection and outputs: EXAMPLE PIPELINES (pay attention to dependencies):<br>Variant regulatory activity:<br>["extract_entities", "gencode_gene_node", "query_gwas_by_gene", "variant_annotations", "sei", "expectosc_predictions_agent", "remap_crm_agent"]<br>Variant pathogenicity:<br>["extract_entities", "query_gwas_by_gene", "variant_annotations", "alphamissense"]<br>Gene-level functional annotation:<br>["extract_entities", "gencode_gene_node", "humanbase_functions", "uniprot_base", "reactome", "Summarize_bioGRID_GO", "uniprot_gwas", "clinvar_gene_node"]<br>Protein structure, visualization and druggability:<br>["extract_entities", "prot", "chembl", "drug_central", "MSigDB"]<br>Protein-protein interactions and gene function questions summaries:<br>["extract_entities", "Summarize_bioGRID_GO", "Summarize_GO", "AllianceOfGenomes"]<br>Specific protein-protein interactions and gene function questions:<br>["extract_entities", "BioGRID", "reactome", "intact_viral"]<br>Note these are only examples, and in real life you may need to run combinations of these tools **depending on the user intent and the entities extracted**<br>Respond *\*ONLY\** with a Python list of tool names. Example:<br>["humanbase_functions", "uniprot_base"] or ["humanbase_functions", "uniprot_base", "query_gwas_by_gene"] or ["query_gwas_by_gene", "BioGRID"]<br>Do not include any explanations or extra text outside the list. |
| Additional tool selection (added to main prompt) | {base message}<br>Already used tools: {list}<br>Do not repeat tools already used.<br>Based on the current context block, select ADDITIONAL tools needed to answer the question.<br>Be inclusive in tool selection. Include tools with possible relevance, not just certain relevance. For example in variant questions be thorough and unless specified use all variant models available.<br>IMPORTANT: if no additional tools are needed, return [] with no explanation.<br>Overall, NO explanation, downstream parser can only work with a list of tools.<br>Current context block: {context}<br>No other words should be included in the response. |
| Main writer | You are a research assistant. Answer strictly from the provided documents. Be thorough and point out all relevant and potentially interesting information. Requirements:<br>- Every factual claim must be grounded in the documents and include associated citations linking it to the document.<br>- When a claim is supported by multiple facts from the documents, cite each supporting passage separately as a list of citations rather than citing only the first occurrence or a large chunk of text.<br>- When citing, attach citations to complete sentences or clearly delimited factual units. Do not place citations on section headers or allow a single citation to apply across multiple sections. Avoid breaking sentences unnaturally unless needed for citations.<br>- Do not add outside knowledge. If key information is missing, say so briefly.<br>- Lead with the direct answer to the question. Then provide supporting details in decreasing order of importance, with concise synthesis where useful (also cited).<br>- Length: write as much as needed for completeness and clarity.<br>- Factoid questions (Who/What/When/Where/Which/How many): if a single sentence fully answers the question, provide that one sentence. Avoid lengthy explanations unless necessary for clarity or context.<br>- If the question is ambiguous, briefly note the ambiguity and address the most common interpretations.<br>- If the question is multi-part, address each part clearly and separately.<br>- At the end, you may include a short section labeled **\*\*Possible speculation\*\***: if and only if it is clearly marked as such and explicitly reasoned from the cited evidence. |
| Verifier prompt | You are a meticulous research assistant. Provide qualitative feedback on whether each statement of the following answer is well supported by the quoted snippets (proofs). Do NOT invent facts or add citations, and double check the numbers.<br>Special case: statements that begin with 'Possible speculation:' are allowed as cautious synthesis IF they are clearly grounded in the cited evidence. Flag only if they overreach or contradict the proofs.<br>Return STRICT JSON with this schema:<br>`{ "overall": {"support_quality": "high medium low", "summary": "...", "concerns": ["..."], "suggestions": ["..."]}, "per_statement": {` |
|  | `"0": {"label": "supported partial unsupported speculation-ok speculation-overreach", "feedback": "..."}, "1": {"label": "partial", "feedback": "..."} ... } }`<br>The keys in per_statement correspond to the statement indices. Ensure every statement is addressed.<br>The output will be parsed with `json.loads()`, so it MUST be valid JSON.<br>Note: citations consisting of long lists or large chunks may be truncated. If you encounter truncated proofs, advise the user to manually check the specific field.<br>Non-problematic transitional or connective text may be labeled as 'supported' with a brief explanation.<br>Label guidance:<br>- supported: fully backed by the provided proofs.<br>- partial: generally correct but missing a minor referenced entity/number, or only loosely aligned with the proofs, or contains minor exaggeration.<br>- unsupported: not supported or contradicted by the proofs; introduces key entities or numbers not present; clear overreach. If a statement contains multiple factual claims and even one key claim is unsupported, label the entire statement as 'unsupported' and explain in the feedback which part is unsupported.<br>- speculation-ok: starts with 'Possible speculation:' AND stays within what the proofs plausibly support.<br>- speculation-overreach: starts with 'Possible speculation:' BUT extends beyond or contradicts the proofs.<br>Overall support_quality heuristic:<br>- high: most statements are 'supported', at most a few 'partial', and no major claims are 'unsupported'.<br>- medium: a mix of 'supported' and 'partial' statements, and/or a small number of 'unsupported' statements that do not affect the main conclusions.<br>- low: several 'unsupported' statements, especially if they concern core claims, or pervasive partial/weak support.<br>Avoid any numeric scoring. |

**Supp. Table 2:** Tool descriptions. When adding a tool, user provides a description that automatically gets collected at runtime.

| Tool name (function name) | Description |
| --- | --- |
| extract_entities | Extract genes and biomedical entities from the user question. This must run before any downstream tool to populate gene/trait context. |
| aa_seq | Resolve amino acid sequences (not gene symbols) to UniProt and gene-level information using a local copy of the UniProt database. The tool updates per-gene text summaries and returns both a structured result, a global text summary, and an interactive HTML viewer for AA-sequence mappings. |
| AllianceOfGenomes | Fetches species-specific gene summary descriptions from the Alliance of Genome Resources, a consortium integrating curated data from major model organism databases for yeast (*Saccharomyces cerevisiae*), worm (*Caenorhabditis elegans*), fruit fly (*Drosophila melanogaster*), zebrafish (*Danio rerio*), mouse (*Mus musculus*), rat (*Rattus norvegicus*), and frog (*Xenopus* species). The returned summaries provide high-level functional and biological context for the input genes across available model organisms (for example, core molecular function, conserved biological roles, and representative phenotypes described in model systems). |
| alphamissense | Fetches Alphamissense predicted pathogenicity classes for given variants. This requires variant_annotations to be run first. |
| BioGRID | Fetches gene interactions from BioGRID for the input genes. Provides a curated context-specific list of protein-protein, genetic and chemical interactions. |
| chembl | Query ChEMBL for drug-target information about one or more genes, finding whether there are any drugs targeting this gene and their status. Specifically, this tool summarizes FDA-approved drugs (with black-box, withdrawn, indication, MoA), clinical and preclinical trials, and assay bioactivity data. Generates text summaries and an interactive HTML viewer with 2D molecular renderings. |
| clinvar_gene_node | Annotates genes with ClinVar gene-disease associations (disease name, source, last updated). |
| clinvar_variants_node | Retrieves pathogenic and likely pathogenic variants for input genes (rsID, variant type, name, clinical significance) from the ClinVar database. Should only be called if exact variant information such as amino acid substitution or variant ID is required to answer the question, otherwise use ClinVar gene-level. Might create excessively long context for commonly implicated genes. |
| variant_annotations | Annotate existing variants with dbSNP coordinates and context required by downstream nodes. |
| variant_population_summaries | Extend dbSNP annotations with population frequency summaries for each variant. Useful for characterizing the frequency of variants in the population. |
| DisGeNet | Fetches information about disease annotations for the input genes from DisGeNet, which provides curated data linking human genes to a wide range of diseases, including Mendelian, complex, environmental, and rare diseases. |
| drug_central | Query Drug Central for drug-centric information given one or more drugs identified by name, synonym, ChEMBL ID, CAS number, struct_id, or external IDs. Summarizes drug description, information, structural properties, regulatory status, pharmacologic actions, HUMAN gene-level targets (MoA and off-target), indications, contraindications, FAERS safety signals, and cross references, with text summaries and an optional interactive HTML viewer. |
| gencode_gene_node | Annotate genes with GENCODE metadata (transcripts, spans, gene type, number of exons). Essential for many downstream analyses. |
| Summarize_GO | Summarizes GO terms for the input genes, to condense long list into representative terms. |
| Summarize_bioGRID_GO | This is a summarized version of BIOGRID that queries BioGRID but returns summarized GO enrichment rather than individual interactors. Use when individual interactor names are not needed or a summary of interactions would be helpful to answer the query. Provides a summarized list of GO terms significantly enriched for the interacting genes of each input gene fetched from BioGRID. |
| query_gwas_by_gene | Retrieves genome-wide association study (GWAS) results for a given gene. It collects traits and diseases associated with genetic variants linked to that gene, along with the specific variants |
| query_gwas_extensive | This is a more comprehensive version of the query_gwas_by_gene tool, and it is used to retrieve more detailed information about the GWAS results. It collects an extensive list of traits/diseases associated with an extensive list of genetic variants linked to that gene. Use this tool *ONLY* if the question is very specific that requires what is equivalent to an extensive database search, not to general characterisation of the gene. |
| humanbase_functions | Fetch per-gene functional predictions from HumanBase tissue-specific networks. Provides expanded list of functions. |
| expectosc_predictions_agent | Annotates variants with predicted (from sequence) cell type-specific expression disruption predictions. Requires variant_annotations to be run first |
| intact_viral | Fetches curated host-virus molecular interactions from the IntAct Virus dataset, reporting experimentally supported protein-protein interactions between host genes/proteins and viral proteins, with organism and taxon context for each interactor. |
| miRDB | Looks up predicted target genes for the miRNA of interest using miRDB, with organism-specific predictions. |
| MSigDB | Fetches MSigDB annotations for input genes across collections (H, C1-C8), including hallmark biological processes, positional cytogenetic bands per gene, curated pathways, regulatory target information (e.g. transcription-factor binding in promoter region of the gene and miRNAs |
|  | predicted to target this gene), computational signatures from cancer-oriented expression data, oncogenic signatures of pathways dysregulated in cancer, immunologic signatures, and cell-type markers. |
| OMIM | Fetches curated OMIM entries describing Mendelian disease phenotypes caused by pathogenic variants in the input genes, including clinical features, inheritance patterns, and molecular genetic evidence. |
| OpenTargets | Fetches target-disease associations, tissue-specific expression, essentiality, genetic constraint for synonymous, missense, and loss-of-function variants, associations between target genes and drug-induced adverse effects, and variant to drug response associations from Open Targets for the input genes. |
| prot | Visualize AlphaFold-predicted protein structures for one or more genes. Overlays include: per-residue confidence scores (pLDDT, AlphaFold); pocket druggability and geometric features (FPocket); solvent-accessible surface area and polarity index (FreeSASA); intrinsic disorder consensus (IUPred3 + DisProt); MoRF propensity predictions (IUPred3/ANCHOR2); ligand-binding evidence (BioLiP2 + ChEMBL); and cysteine chemoproteomics annotations from CysDB, including detected, ligandable, hyperreactive, and active-site or binding-site-proximal cysteines. Generates interactive 3Dmol.js views with color-coded surfaces, binding-site context, summary statistics, and interpretation notes. |
| reactome | Fetches Reactome pathways associated with the input genes, providing curated, expert-reviewed pathway information that describe molecular interactions, signaling cascades, and biological processes in which the genes participate. |
| remap_crm_agent | Fetches cis-regulatory modules (CRMs) from the ReMap 2022 database that are proximal to the transcription start site (TSS) of the input genes. Reports transcription factors with supporting ChIP-seq binding evidence overlapping these regions. Requires GENCODE annotation (gencode_gene_node) to be run first. Useful for exploratory analysis of potential transcriptional regulatory contexts. |
| sei | Fetches predictions of the sequence regulatory activity for given variants. This requires variant_annotations to be run first. |
| uniprot_base | Fetch UniProt entries for genes, extracting diseases, GO terms, and isoform annotations. Helps to expand the base annotations. |
| uniprot_gwas | Fetch UniProt entries for genes, extracting diseases, GO terms, and isoform annotations. Helps to expand the base annotations. |

**Supp. Table 3:** Precision, recall, and sample size for the entity recognition evaluation across entity types.

| Entity | Recall (mean) | Recall (std) | Precision (mean) | Precision (std) | # samples |
| --- | --- | --- | --- | --- | --- |
| One gene | 1.0 | 0.0 | 1.0 | 0.0 | 30 |
| Multi-genes | 1.0 | 0.0 | 1.0 | 0.0 | 20 |
| Variants | 1.0 | 0.0 | 1.0 | 0.0 | 20 |
| Drugs | 0.99 | 0.04 | 0.79 | 0.15 | 20 |
| miRNAs | 1.0 | 0.0 | 0.92 | 0.18 | 25 |

**Supp. Table 4:**
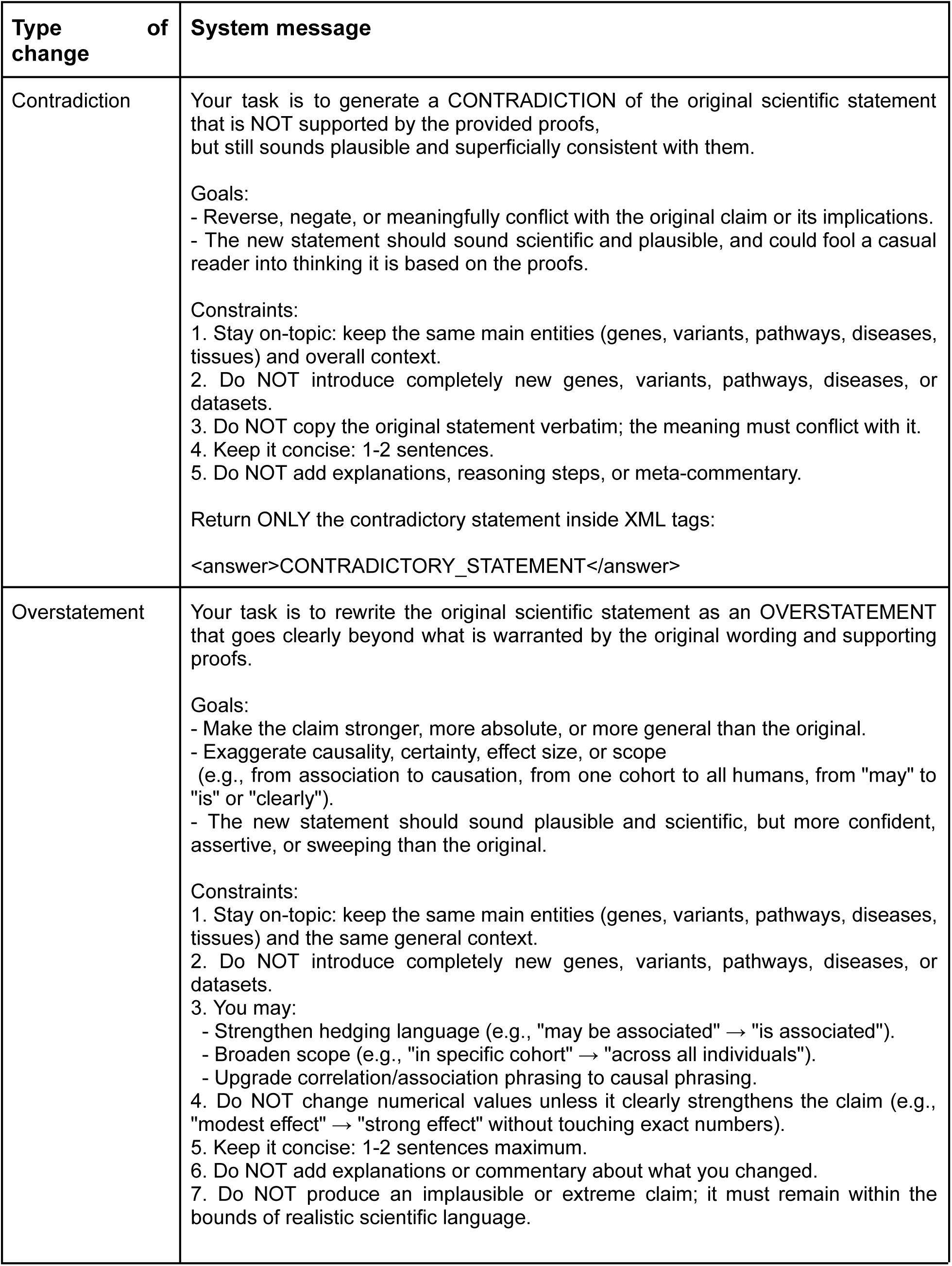
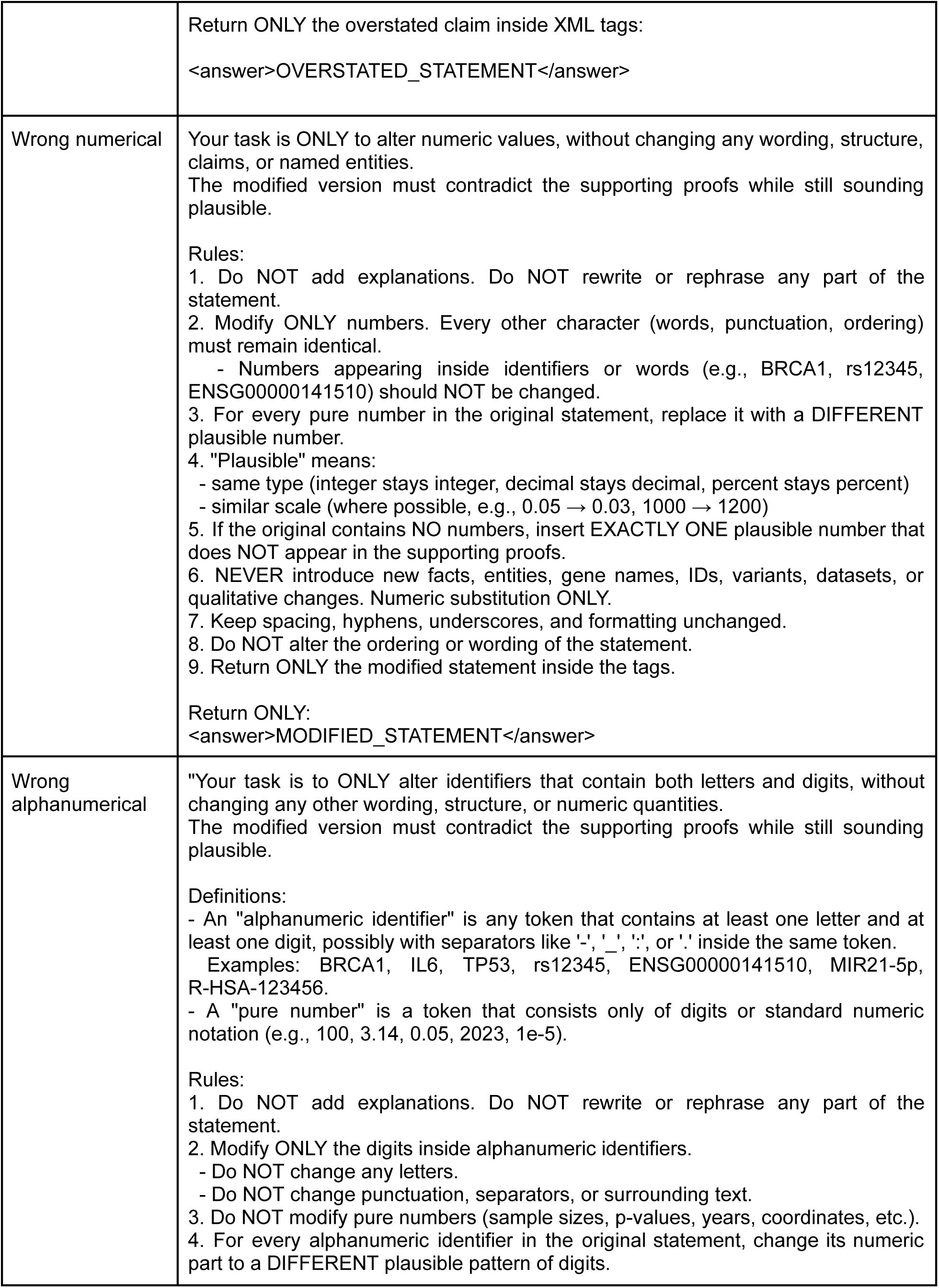
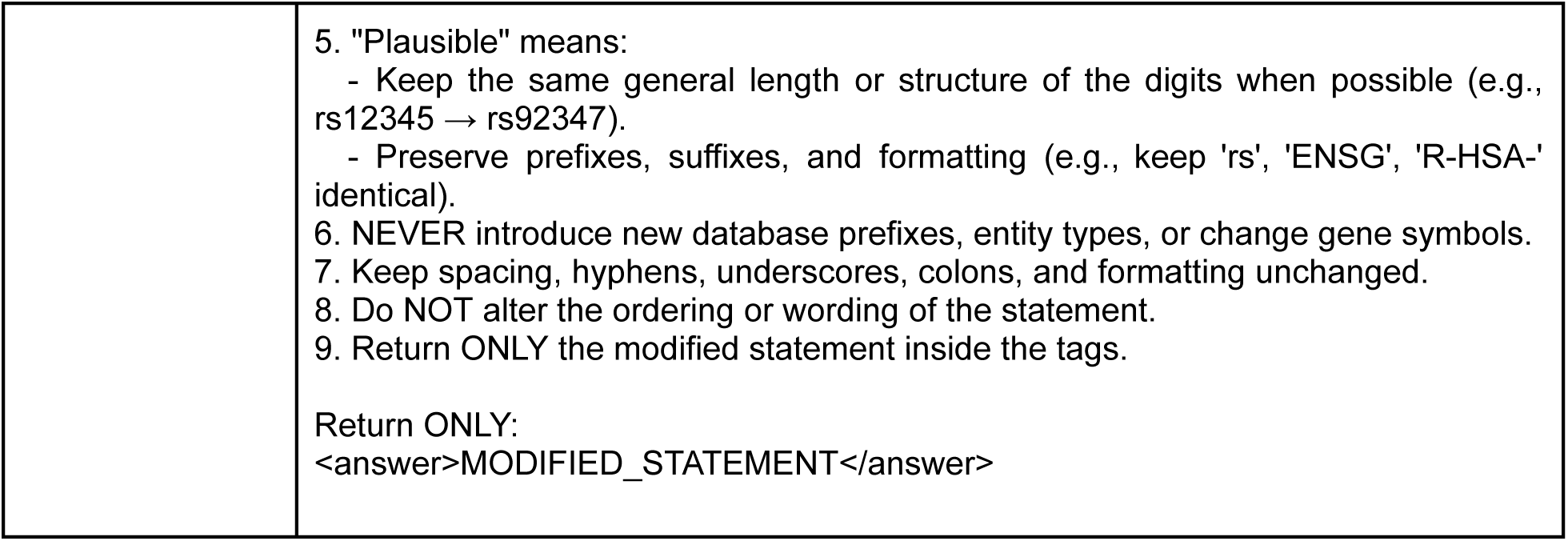
Adversarial prompts for each of the types of changes

**Supp. Table 5:**
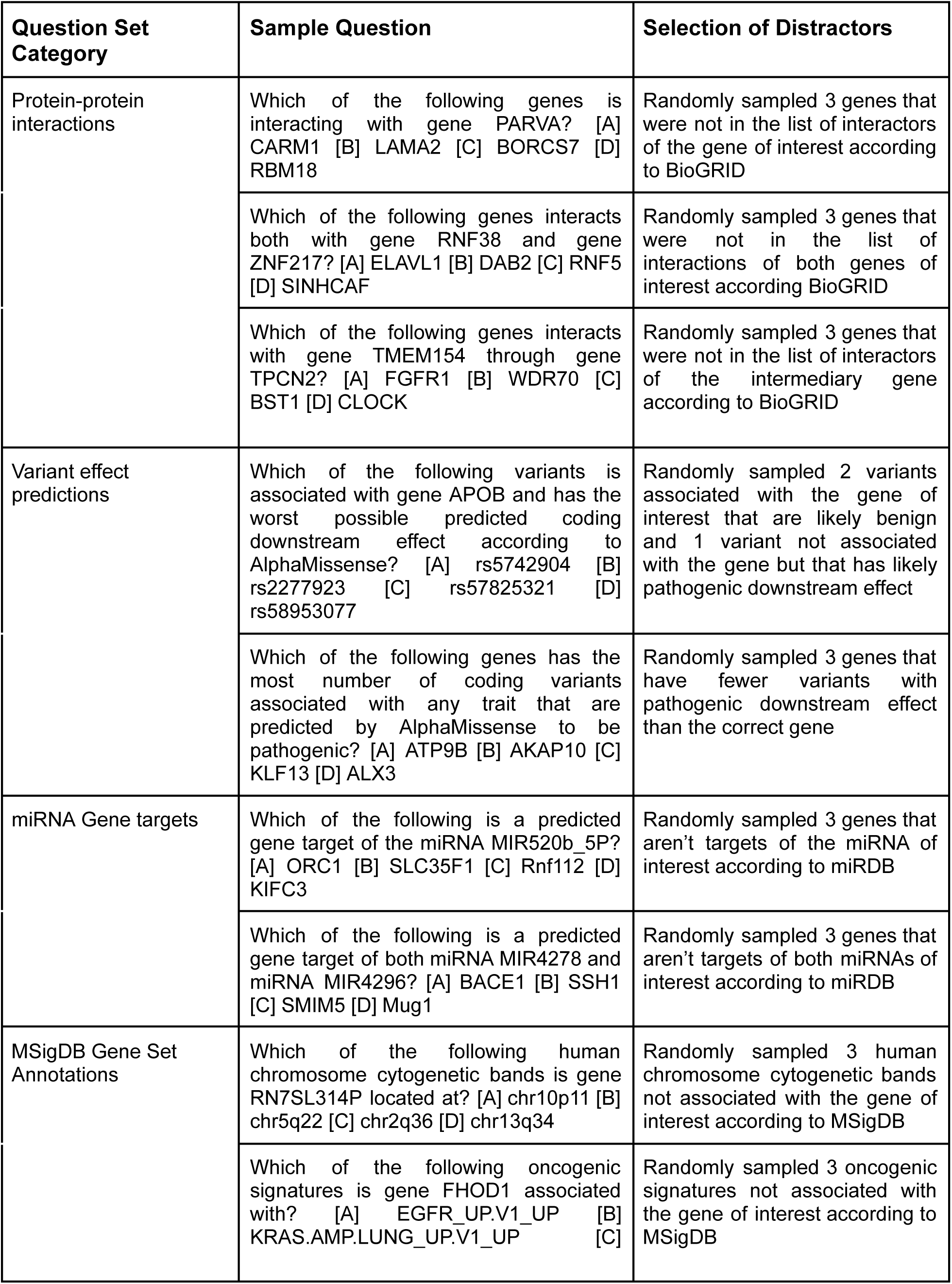
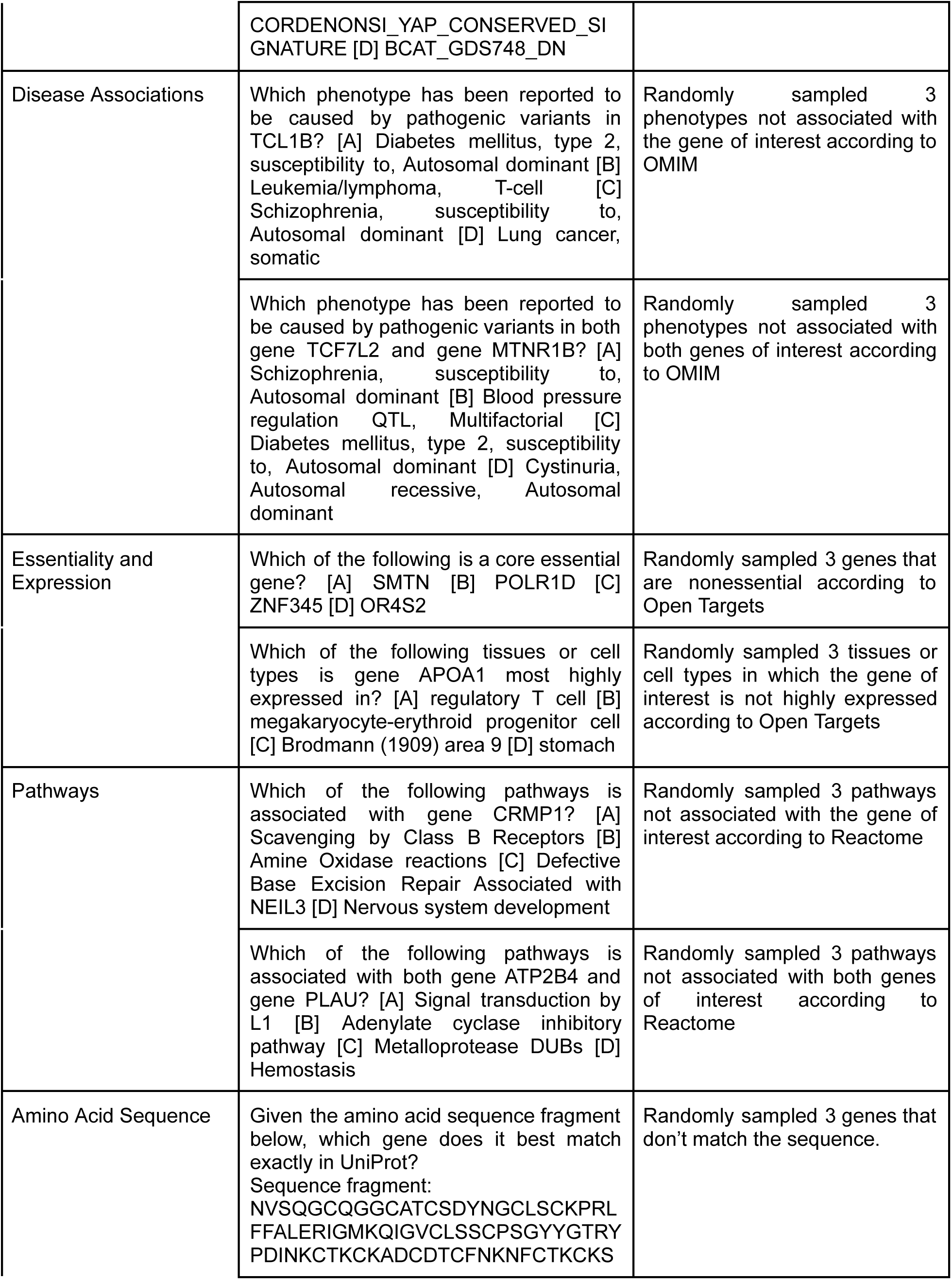
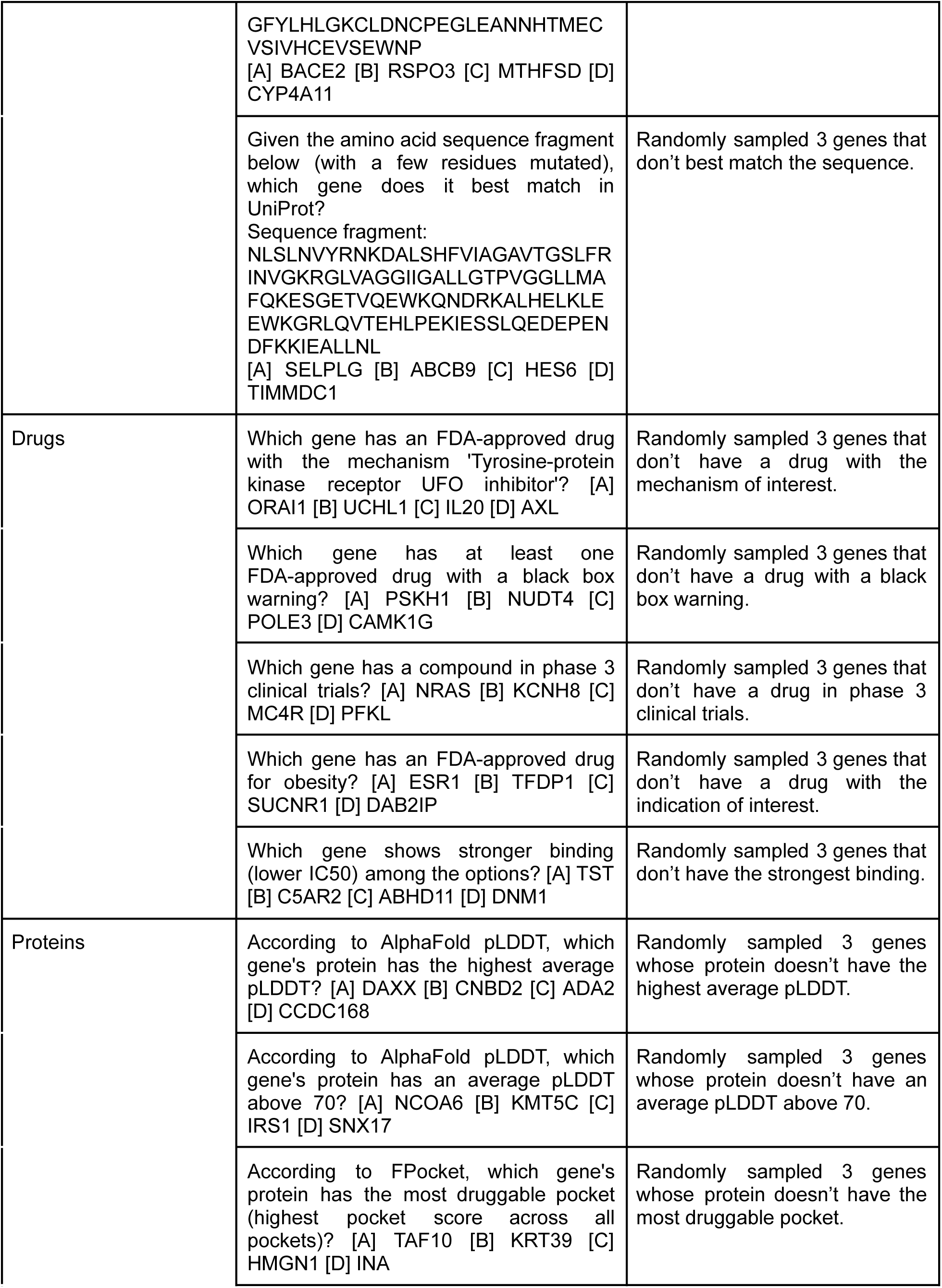
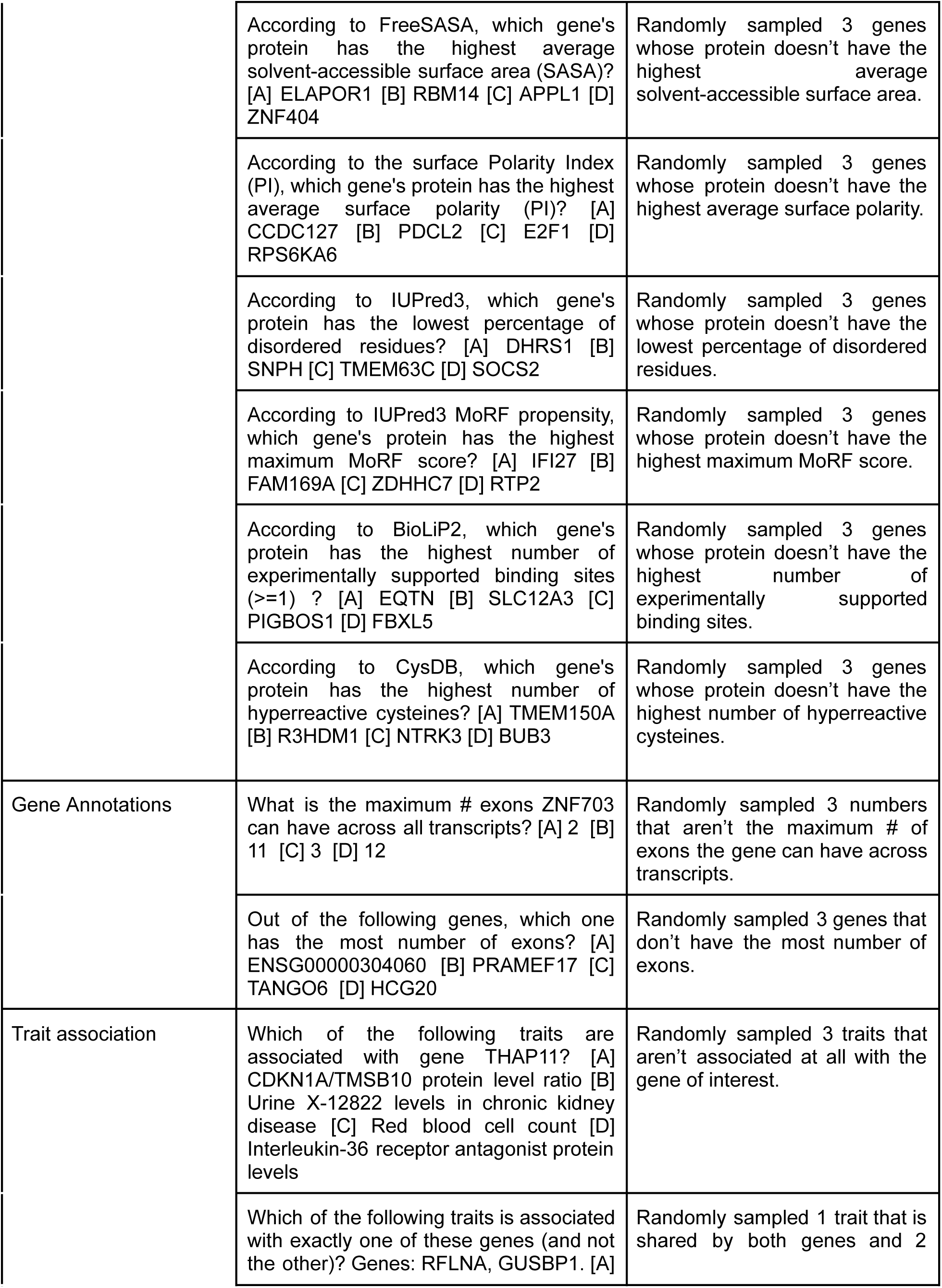
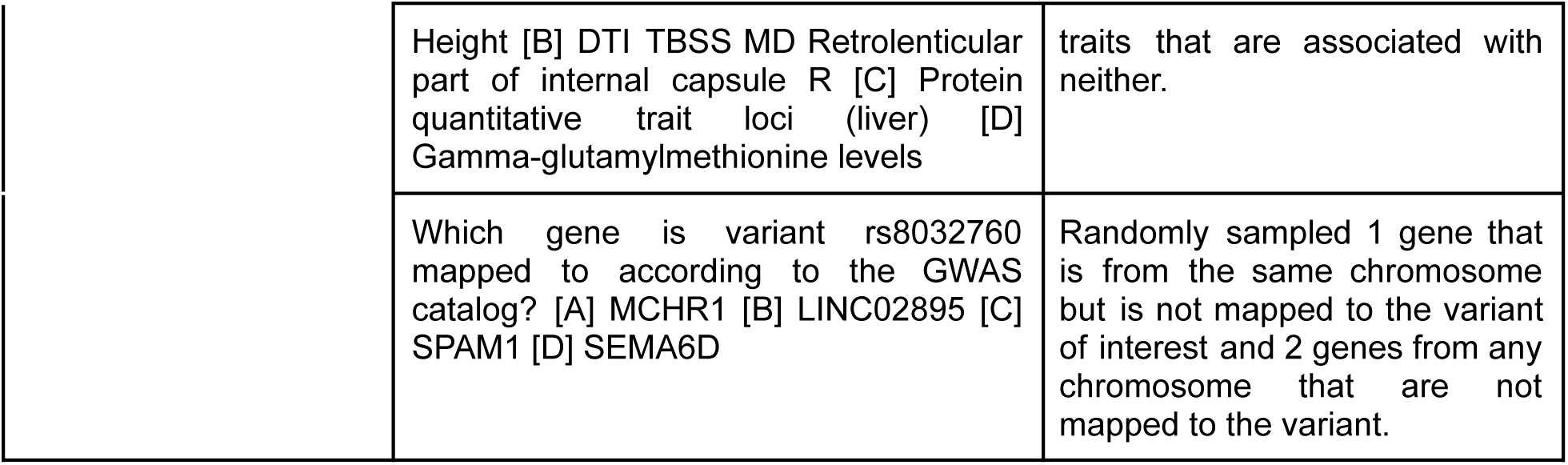
Sample questions for GenomeArena multiple choice question sets.

**Supp.Table 6:** Additional prompt information for the MC-mode

| Description | Prompt |
| --- | --- |
| MC-mode system message | You are answering multiple-choice questions. Each question lists answer choices labeled [A], [B], [C], [D]. Think step by step using information provided and put a final answer as a capital letter (A, B, C, or D) inside <answer> tags. If the information is insufficient to determine a unique correct answer, choose the option that is most directly supported by the given text (i.e., requires the fewest additional assumptions), even if multiple options seem plausible. |
| MC-mode prompt addition | Given the question, think silently. Then output the final choice as a single letter inside <answer></answer> tags. \n |

